# ROR2 drives right ventricular heart failure via disruption of proteostasis

**DOI:** 10.1101/2025.02.01.635961

**Authors:** Hali Hartman, Genevieve Uy, Keita Uchida, Emily A. Scarborough, Yifan Yang, Eric Barr, Spencer Williams, Ryan T. Santilli, Sanam L. Kavari, Jeff Brandimarto, Li Li, Ling Lai, Joanna Griffin, Nora Yucel, Swapnil Shewale, Hari Rajagopal, Deborah M. Eaton, Tanis Dorwart, Kenneth C. Bedi, Crystal S. Conn, Kenneth B. Margulies, Benjamin Prosser, Zoltan Arany, Jonathan J. Edwards

**Affiliations:** Cardiovascular Institute, Children’s Hospital of Philadelphia, Philadelphia, PA, USA; Department of Physiology, Pennsylvania Muscle Institute, Perelman School of Medicine, University of Pennsylvania, Philadelphia, PA, USA; Cardiovascular Institute, Perelman School of Medicine, University of Pennsylvania, Philadelphia, PA, USA; Department of Radiation Oncology, Perelman School of Medicine, University of Pennsylvania, Philadelphia, PA, USA

## Abstract

**Background:** No therapies exist to reverse right ventricular failure (RVF), and the molecular mechanisms that drive RVF remain under studied. We recently reported that the developmentally restricted noncanonical WNT receptor ROR2 is upregulated in human RVF in proportion to severity of disease. Here we test the mechanistic role of ROR2 in RVF pathogenesis.

**Methods:** ROR2 was overexpressed or knocked down in neonatal rat ventricular myocytes (NRVMs) and then characterized using confocal microscopy, RNAseq, proteomics, proteostatic functional assays, and pacing to assess contractile properties. The impact of cardiac ROR2 expression was evaluated in mice by AAV9-mediated overexpression and by AAV9-mediated delivery of shRNA to knockdown ROR2 in a pulmonary artery banded pressure overload model of RVF. ROR2-modified mice were evaluated by echocardiography, histology, and RV protein synthesis and proteasome capacity.

**Results:** In NRVMs, we find that ROR2 profoundly dysregulates the coordination between protein translation and folding. This imbalance leads to excess protein clearance by the ubiquitin proteasome system (UPS) with dramatic impacts on sarcomere and cytoskeletal structure and function. Inhibiting the UPS or restoring chaperone expression is sufficient to partially rescue ROR2-induced structural and contractile deficits in cardiomyocytes. In mice, forced cardiac ROR2 expression is sufficient to disrupt proteostasis and drive RVF, while conversely ROR2 knockdown partially rescues proteostasis and RV structure and function in a pressure overload model of RVF.

**Conclusions:** In sum, ROR2 is a key driver of RVF pathogenesis through proteostatic disruption and, thus, provides a promising target to treat RVF.

## Introduction

Right ventricular failure (RVF) is a widespread problem for which there are no proven therapies. Nearly half of patients with left ventricular failure (LVF) develop RVF, and RVF doubles their risk of death.^1,2^ RVF is also the leading cause of hospital admission and death in pulmonary hypertension.^3–5^ For LVF, there are several therapies that can reduce patient morbidity and mortality.^6^ In contrast, no therapies have proven efficacious for patients with RVF, and in fact, some medications that benefit LVF led to worse outcomes.^7–13^ RVF thus fundamentally differs from LVF, and finding new therapies for RVF will require a deeper understanding of targetable molecular drivers of RVF.^14,15^

We recently analyzed a large cohort of human hearts from patients transplanted for dilated or ischemic cardiomyopathy, and discovered that the noncanonical WNT receptor ROR2—usually developmentally restricted—is robustly upregulated in failing RVs, and its expression correlated with RVF severity.^16^ ROR2 is essential for the development of multiple organ systems, including the heart.^17^ *Ror2* knockout caused ventricular septal defects in mice, while loss of *Ror1* did not impact heart development.^17–19^ However, combined *Ror1/Ror2* double knockout caused more severe developmental pathologies including complex conotruncal defects and ventricular noncompaction. The mechanistic role of ROR2 in RVF, however, has not been explored.

We investigated whether ROR2 induction contributes to RVF and identified a previously unrecognized role for ROR2 in regulating cardiomyocyte proteostasis. ROR2 induction disrupts the balance between protein synthesis and folding capacity and increases ubiquitin proteasome system (UPS)-mediated protein degradation. This UPS response is either inadequate or excessive to the point being maladaptive, further contributing to RVF pathogenesis. In a mouse model of RVF, preventing ROR2 induction ameliorated both RV proteostatic disruption and RVF. These findings indicate that targeting ROR2-driven proteostatic disruption may offer a new therapeutic approach for RVF.

## Methods

Detailed methods are provided in the Supplemental Material. Source data supporting the findings and displayed in main figures and supplemental figures are also included in the Supplemental Material. RNAsequencing data generated for this study was deposited in the Gene Expression Omnibus under the accession number GSE278627. The mass spectrometry proteomics data have been deposited to the ProteomeXchange Consortium via the PRIDE partner repository with the dataset identifier PXD05655.^20^

### Animal Studies

All animal work was performed under University of Pennsylvania Institutional Animal Care and Use Committee approved protocols (#805255, #805309, and #807751). Neonatal rat ventricular myocytes (NRVMs) were isolated from 1-2 day old Sprague Dawley rats after obtaining pregnant rats from Charles River Laboratory and maintained in culture as previously described.^21^ C57BL/6J mice obtained from Jackson Laboratory were used for all mouse experiments studies.

### Human Studies

Procurement of all human RV myocardial tissue was performed in accordance with protocols approved by the University of Pennsylvania Institutional Review Board (approval 802781) and the Gift-of-Life Donor Program, Philadelphia, PA. Prospective informed consent for research use of surgically explanted tissue was obtained from the transplant recipients or from the next-of-kin in the case of deceased organ donors, as previously described.^22^

### Statistical Analyses

RNAseq and proteomics data were analyzed in RStudio using the voom function from the Limma package (version 3.56.1) to transform and normalize the data, and the toptable function to output the differential analysis result. Two group and three group statistical comparisons were performed in GraphPad Prism v10 using T-test/Mann Whitney U or One-Way ANOVA/Kruskal-Wallis/Two-Way ANOVA based on normality and number of independent variables.

## Results

### Cardiomyocyte ROR2 expression is developmentally restricted and reactivated in RV failure

ROR2 is necessary for heart development and its expression is broadly absent in healthy adult tissues, but the temporal nature for its loss and its histologic pattern in the early postnatal heart has not been well characterized.^23^ By qPCR and western blot, we observe maturity dependent loss of ROR2 with about 70% reduction in ventricular ROR2 from P2 to P7 and near loss of ROR2 by P21 (**Fig. 1a, Supplementary Fig. 1a/b**). Myocardial ROR2 at birth is observed predominantly in cardiomyocytes with localization to the cell membrane and nucleus (**Fig 1b** and **Supplementary Fig. 1c**). With postnatal maturation, ROR2 is minimally retained and mostly observed in the atria and subepicardium (**Fig 1a** and **Supplementary Fig. 1c/d**), consistent with prior single nuclear RNAseq datasets.^24,25^ Nuclear localizing ROR2 has been observed in some cancers, which has been linked to worse outcomes. Therefore, we further evaluated ROR2 localization using a protein fractionation method that separates nuclear depleted cytoplasmic proteins (“NCD”) from a nuclear enriched fraction that also contains cytoskeleton proteins (“NCE”) (**Supplementary Fig. 1e/f**).^26–28^ We found most ROR2 localized to the NCD fraction, but the temporal loss of ROR2 is comparable across total, NCD, and NCE fractions.

**Fig. 1.**
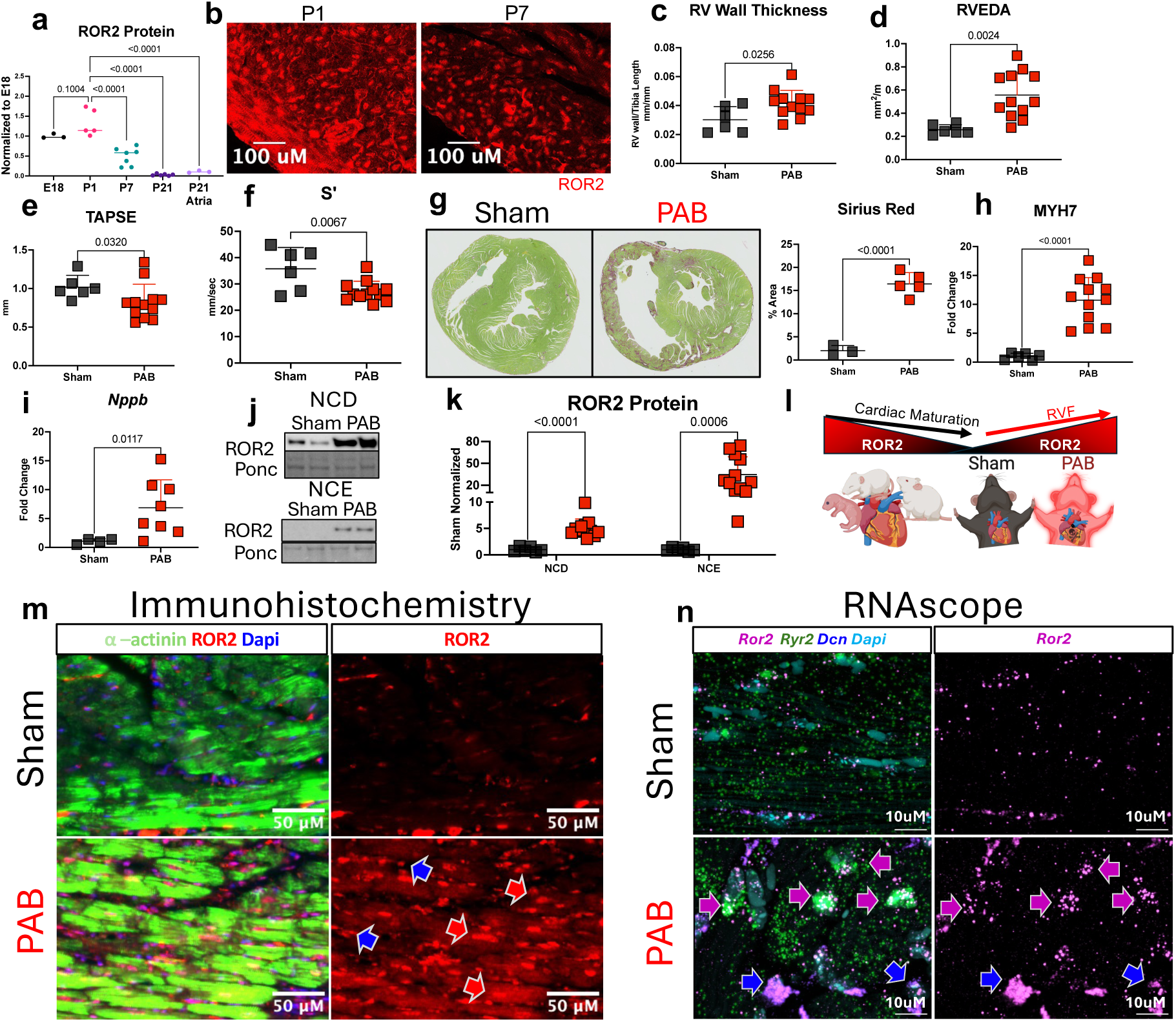
ROR2 is a developmentally restricted protein that is robustly upregulated in RVF. (**a**) ROR2 protein expression was measured in the ventricular tissue using thiourea extraction by western from P2 to P21 and in P21 atria (N=3-7/age group; statistics by One-Way ANOVA with Two step FDR correction and T-test for P21 ventricle vs atria). (**b**) ROR2 immunohistochemistry was performed to evaluate localization, which is observed at the cell membrane and nucleus. (**c-f**) PAB was performed in 9-week-old C57BL/6J male mice with echocardiogram performed after 2 weeks to assess RV remodeling and systolic function, Sirius red staining to assess for fibrosis quantified using ImageJ **(g,** 10X magnification), western blot to assess induction of fetal MYH7 (**h**) and qPCR to assess expression *Nppb* in the RV free wall. Protein fractionation was performed to assess expression of ROR2 in the NCD (cytoplasmic, triton soluble) and the NCE fractions (triton-insoluble) (**j-l**). ROR2 immunohistochemistry (**m**, 20X magnification) and RNAscope (**n**, 63X magnification), was performed to assess cell type pattern of expression for ROR2 induction in PAB using ⍺-actinin and *Ryr2* to identify cardiomyocytes and *Dcn* to identify fibroblasts in PAB RV. Statistics for PAB analyses by unpaired t-test or Mann-Whitney, depending on normality (n= 3-6 for sham, n=5-12 for PAB).

We previously found that ROR2 is re-expressed in human RVF in a severity dependent manner, localizing to both cardiomyocytes and fibroblasts.^16^ This pattern differs from a prior study in mouse pressure overload LVF where ROR2 induction was restricted to noncardiomyocytes, so we next tested if and where ROR2 is reactivated in the failing mouse RV.^29^ To that end, we used a pulmonary artery band (PAB) pressure overload model to generate RVF. With an average pressure gradient of ∼60-70mmHg, PAB mice developed clear markers of pathologic RV remodeling and RVF with increased RV mass, and echocardiographic RV hypertrophy, dilation, systolic dysfunction, and tricuspid insufficiency (**Fig. 1c-f** and **Supplementary Table 1**). PAB also caused RV wall thickening and diffuse RV fibrosis by histology (**Fig. 1g**). PAB RVs also demonstrate increased expression of the classic heart failure markers fetal myosin heavy chain protein MYH7 and the natriuretic peptide transcript *Nppb* (**Fig. 1h/i**). In parallel, ROR2 was induced ∼5-fold in the NCD fraction and more than 30-fold in the NCE fraction. (**Fig. 1j/k**).

Thus, the normally developmentally restricted ROR2 is robustly reexpressed in mouse RVF (**Fig 1l**). Interestingly, NCE ROR2 strongly correlated to RV functional and structural changes in PAB including the severity of RV dilation (r=0.66), TAPSE (r= -0.69), and fractional area change (r= - 0.60) (all p<0.05, **Supplementary Table 1**). In contrast, NCD ROR2 showed no significant correlation to any of these RV functional or structural parameters. Induced ROR2 localized to both cardiomyocytes and fibroblasts in the PAB RV by immunohistochemistry and RNAscope.

ROR2 protein in cardiomyocytes localized to the cytoplasm, membrane, and nucleus (**Fig. 1m**) similar to ∼P2-P7 mouse RV and consistent with the NCD/NCE fractionation. This was also consistent with our previous findings in human RVF immunohistochemistry and further supported by complementary immunohistochemistry and RNAscope of human RVs in this study (**Supplementary Fig. 2a/b).** We conclude that ROR2 expression, particularly the NCE fraction, is strongly induced in mouse RVF and increases in proportion to RVF severity.

### ROR2 regulates cardiomyocyte morphology

Since little is known about the role of ROR2 in cardiomyocytes, we directly tested the impact of ROR2 induction and loss using NRVMs. We cultured cells on patterned substrates to promote sarcomere alignment, allowing more robust structural characterization. We overexpressed human ROR2 (ROR2^OE^) or knocked down native *Ror2* by shRNA (ROR2^KD^) using adenovirus with a GFP reporter (**Fig. 2a/b**). Adeno-GFP alone was used as a control. ROR2^OE^ and ROR2^KD^ caused dichotomous morphologic changes. ROR2^OE^ NRVMs had fragmented sarcomeres and were significantly smaller, driven by reduction in the longitudinal major axis, with a rounded appearance and smaller aspect ratio (major/minor axis) (**Fig. 2c-e**). In contrast, ROR2^KD^ NRVMs were slightly larger without affecting the aspect ratio. We also observed opposing effects on cell-cell junctions with ROR2^KD^ increasing and ROR2^OE^ decreasing peripheral enrichment of the intercalated disc (ICD) localizing proteins α-actinin, β-catenin, and Cx43 (**Fig. 2f-h** and **Supplementary Fig. 3a**). Total Cx43 and α-actinin staining intensity and protein expression by western blot increased in ROR2^OE^ NRVMs, while β-catenin levels were stable, indicating that reduced peripheral staining represents redistribution of the proteins rather than loss (**Fig 2i-l)**. Thus, ROR2 regulates gross cardiomyocyte morphology and both abundance and peripheral localization of ICD/junction proteins.

**Fig. 2.**
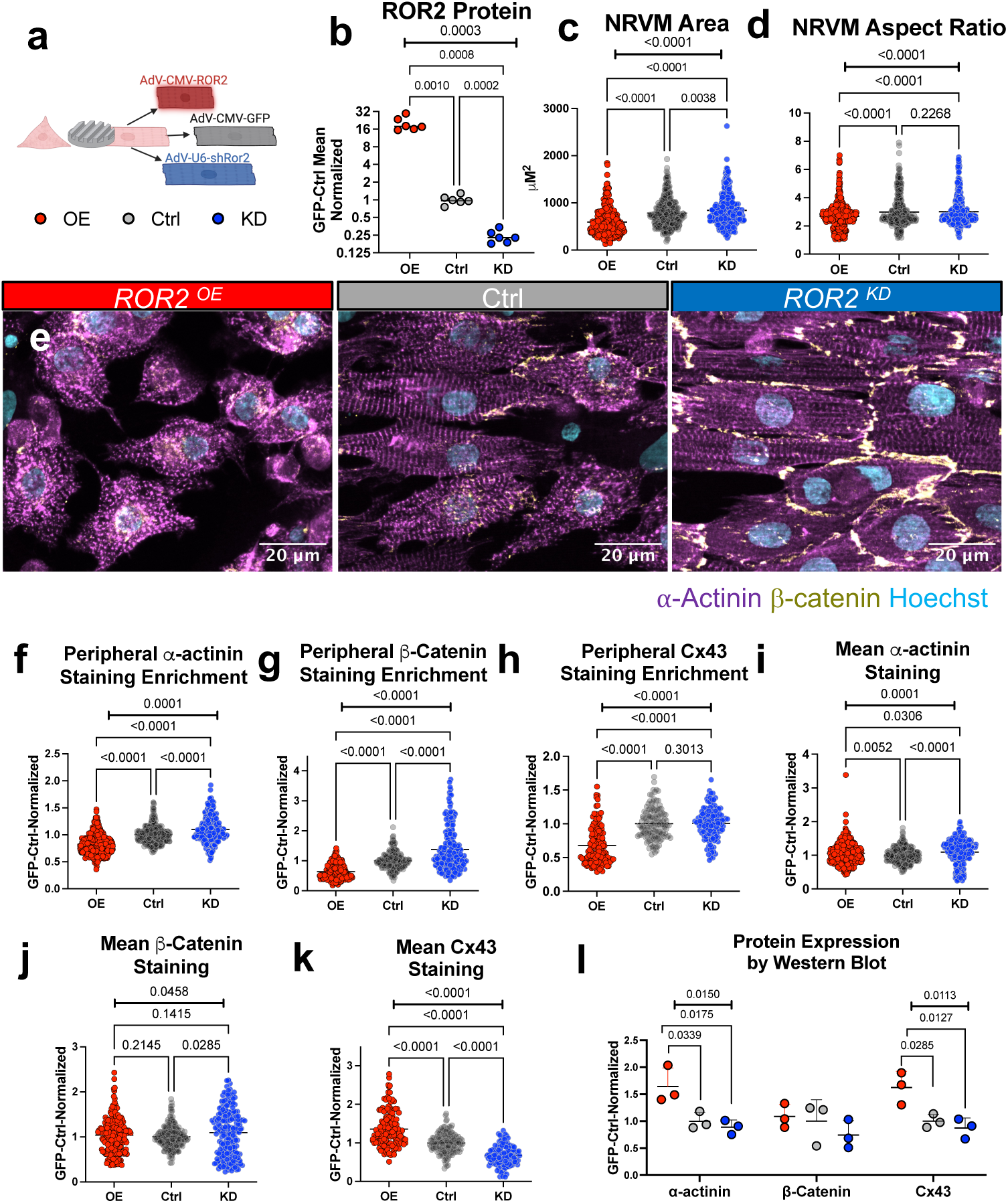
ROR2 regulates in vitro cardiomyocyte morphology. (**a/b**) NRVMs were cultured on nanopatterned surface and received adenovirus to overexpress human ROR2, knockdown ROR2, or express GFP alone. NRVM structure was evaluated by Airyscan confocal microscopy (**c-e**, 63X magnification) using FIJI to generate maximum intensity projections and quantify cell size and aspect ratio (major/minor axis using the fit ellipse). (**f-k**) FIJI was used to quantify mean staining intensity; peripheral intensity enrichment was further measured as described in the methods by subtracting the inner cell integrated intensity (total area shrunken by 2μm) from the full cell integrated for α-actinin, β-catenin, and Cx43 (N=8-9 NRVM isolations, n= 341-411 cardiomyocytes/group). Total protein expression for these were also assessed by western blot (n=3/group). One-Way ANOVA with two-stage step up Benjamini correction).

### ROR2 affects cardiomyocyte ubiquitin-proteasome system, protein folding, and synthesis

We used RNAseq of ROR2^OE^ and ROR2^KD^ NRVMs to investigate potential mechanisms for these NRVM structural phenotypes (**Fig. 3a** and Supplementary Data 4). We observed that β-catenin responsive transcriptional targets were not significantly impacted, indicating no significant role for canonical WNT signaling (**Supplementary Table 2** and **Data 5**). Instead, gene-set enrichment analysis revealed changes in protein turnover gene programs, which included upregulation ubiquitin ligase activity genes and downregulation for ubiquitin-dependent protein catabolism genes in ROR2^OE^ (**Fig. 3b**). Interestingly, at the protein level, total ubiquitin was significantly decreased in ROR2^OE^ but unchanged in ROR2^KD^ (**Fig 3c/d**). We tested whether the reduction of ubiquitinated proteins in ROR2^OE^ NRVMs was driven by increased turnover or decreased ubiquitination by blocking clearance of ubiquitinated proteins using the competitive proteasome inhibitor MG132. As expected, MG132 increased accumulation of total ubiquitin across all groups, with ROR2^OE^ NRVMs demonstrating significantly greater accumulation of proteasome directing K48 ubiquitin (**Fig 3e/f** and **Supplementary Fig 3b**).

**Fig. 3.**
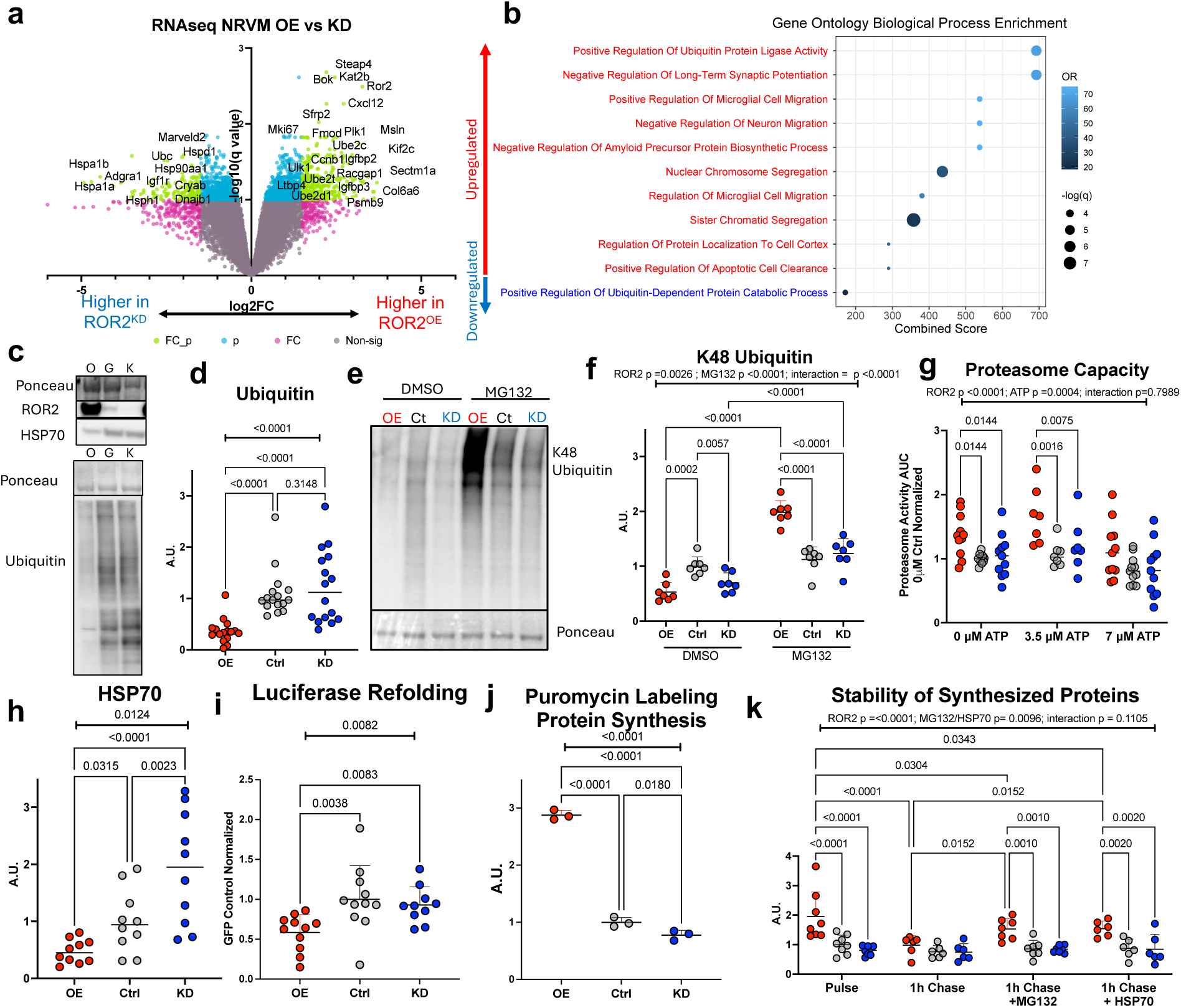
ROR2 regulates in vitro cardiomyocyte proteostasis. RNAseq with Voom/Limma was performed to assess differential gene expression between ROR2^OE^ and ROR2^KD^ NRVMs, with volcano plot (**a**) highlighting notable genes including transcriptional changes in the proteostatic pathways including folding (e.g. *Hspa1a/b, Hsph1, Cryab*) and pathway enrichment analysis (**b**) performed by Gene Ontology Biological Processes (n=4/group). Western blot was used to evaluate total ubiquitin (**c/d**, n=16/group) and HSP70 (**c/h**, n=10/group) between ROR2^OE^, ROR2^KD^, and controls. NRVMs were treated with 1μM MG132 or DMSO for 24 hours prior to collection to assess the flux of K48-linked ubiquitin clearance by the UPS (**e/f**, n=7/group). Proteasome capacity was measured using the chymotrypsin proteasome substrate Suc-LLVY-AMC with cell lysates in the absence or presence of indicated supplemented ATP (**g**, n=7-11/group and ATP condition). Protein refolding was measured using heat shocked luciferase (**i**, n=11/group), measured by luminescence after addition of luciferin. Protein synthesis rates were measured using 0.55 μg/mL tyrosyl-tRNA analogue puromycin for 1 hour pulse with immediate collection (**j**) or with a subsequent 1 hour chase (**k**) including cycloheximide (20 μg/mL) to block further translation with or without MG132 (20 μM) or HSP70^OE^ (n=6-8/group). Statistical comparisons were performed with One-Way ANOVAs or Two-Way ANOVAs as indicated on graphs, with two-stage step up Benjamini correction.

Interestingly, ROR2^KD^ demonstrated lower basal K48 ubiquitin with intact accumulation of both total and K48-linked ubiquitin. We next measured maximal proteasome capacity by incubating cell lysates with Suc-LLVY-AMC—a fluorescent reporter containing a peptide substrate of the chymotrypsin-like proteasome subunit (PSMB5). ROR2^OE^ increased PSMB5 maximal capacity at 0 and 3.5μM ATP, and trended towards an increase at 7μM ATP, without affecting total PSMB5 expression (**Fig 3g** and **Supplementary Fig 3c)**. Whereas, ROR2^KD^ did not impact PSMB5 expression or maximal capacity, mirroring the intact accumulation of ubiquitin. Thus, these data indicate that ROR2 expression significantly alters the UPS, with high ROR2 promoting K48 ubiquitination and proteasome-mediated turnover while low ROR2 decreases K48 ubiquitination without affecting proteasome capacity, indicating additional levels of proteostatic regulation.

In addition to UPS related transcriptional changes, the two HSP70 paralogs *Hspa1a* and *Hspa1b* exhibited the highest magnitude reduction in ROR2^OE^ (log_2_FC <-4.7, adj p =0.059, both). Multiple other chaperones were also trending as significantly downregulated (adj p <0.1) including *Hsph1/Hspa4* (HSP105/110), *Hsp90aa1/ab1*, *Dnajb1/a1 (*HSP40), and *Cryab* (**Fig 3a)**. At the protein level, most tested chaperones including HSP70 were significantly increased in ROR2^KD^. Whereas, in ROR2^OE^, HSP70 was significantly decreased, and other tested chaperones were unchanged (**Fig. 3c/h** and **Supplementary Fig 3d**). To directly measure protein folding capacity, we used a luciferase refolding assay in which heat-shocked recombinant luciferase is allowed to refold in cell lysates supplemented with protease inhibitor and then interrogated by its ability to cleave luciferin thereby generating light (**Fig. 3i**).

Consistent with reduced HSP70 expression, refolding of heat-shocked recombinant luciferase was significantly impaired in ROR2^OE^ lysates. Despite globally increased chaperone expression, luciferase refolding was unaffected in ROR2^KD^. This result may be explained by GFP control lysates exceeding a minimum HSP70 threshold to sufficiently refold heat shocked luciferase or by ROR2^KD^ also increasing HSP105/110, which can lead to premature release of protein substrates that diminishes refolding efficiency (**Supplementary Fig 3c**).^30^

Given the ROR2-responsive effects on protein folding and UPS, we next tested whether other aspects of protein production or stability may also be disrupted. We quantified global protein synthesis rates by using the tyrosyl tRNA analogue puromycin and radiolabeled ^35^S-Met to label newly synthesized proteins. By puromycin labeling, ROR2^OE^ resulted in ∼2-3-fold increase in protein synthesis and ROR2^KD^ modestly decreased translation (**Fig. 3j/k)**. ^35^S-Met labeling revealed similar results with ∼1.5-fold increase in ROR2^OE^ and ∼0.6-fold reduction in ROR2^KD^ (**Supplementary Fig. 3e**). ROR2-driven effects on protein synthesis were unlikely a secondary effect of altered chaperone expression, as HSP70 overexpression (HSP70^OE^) did not reverse the differential translation rates or ubiquitin levels (**Supplementary Fig. 3f/g**). However, given the observed differences in protein synthesis, folding, and proteasome capacity with ROR2 modulation, we considered whether the stability of newly synthesized proteins differed across groups. To that end, we performed a pulse chase experiment consisting of a 1 hour of puromycin labeling followed by a 1hr incubation with cycloheximide to block further protein synthesis with or without MG132 or HSP70^OE^. We found that turnover of newly synthesized proteins was ∼2-fold higher in ROR2^OE^ (∼50% vs. 25%) compared to controls, and ROR2^KD^ demonstrated decreased turnover (<10%) indicating increased protein stability. Strikingly, MG132 and HSP70^OE^ equivalently increased the stability of newly synthesized proteins ROR2^OE^ and control NRVMs, but these only reached statistical significance for ROR2^OE^. We also tested protein stability using the constitutive UPS substrate GFPμ, which required generating ROR2 modified NRVMs lacking a GFP reporter (**Supplementary Fig. 3h-k**). By microscopy, flow cytometry, and western blot, GFPμ was consistently lowest in ROR2^OE^ and highest in ROR2^KD^, even in the presence of MG132. Using bafilomycin, we excluded the possibility that autophagy contributed to GFPμ clearance independently or in combination with proteasome inhibition. Of note, although GFPμ remained lowest in ROR2^OE^, the relative increase of GFPμ by MG132 was most pronounced in ROR2^OE^. Comparing these results with the puromycin pulse chase experiments highlights the importance of considering total protein synthesis rates and baseline GFPμ levels when interpreting GFPμ flux. These results also suggest that ROR2 increases protein synthesis in a biologically selective manner. Taken together, these data show that ROR2 profoundly disrupts cardiomyocyte proteostasis by promoting protein synthesis while simultaneously suppressing folding capacity, priming NRVMs to clear proteins through the UPS.

### ROR2 impairs cardiomyocyte structure and function via Proteostatic Disruption

Next, to determine if changes in the proteostatic balance were responsible for the NRVM morphologic changes observed with ROR2^OE^, we repeated the morphology evaluations with a 24-hour incubation of 1 μM MG132. Proteasome inhibition significantly abrogated the impact of ROR2^OE^ on NRVM morphology with improvement in the sarcomere pattern, aspect ratio, and peripheral localization of α-actinin localization (**Fig. 4a-c** and **Supplementary Fig. 3m**). We also tested the impact of increased folding capacity by HSP70^OE^, which similarly improved ROR2^OE^ NRVM morphology by increasing cell area and aspect ratio, but not to the extent of proteasome inhibition (**Fig. 4d-f**).

**Fig 4.**
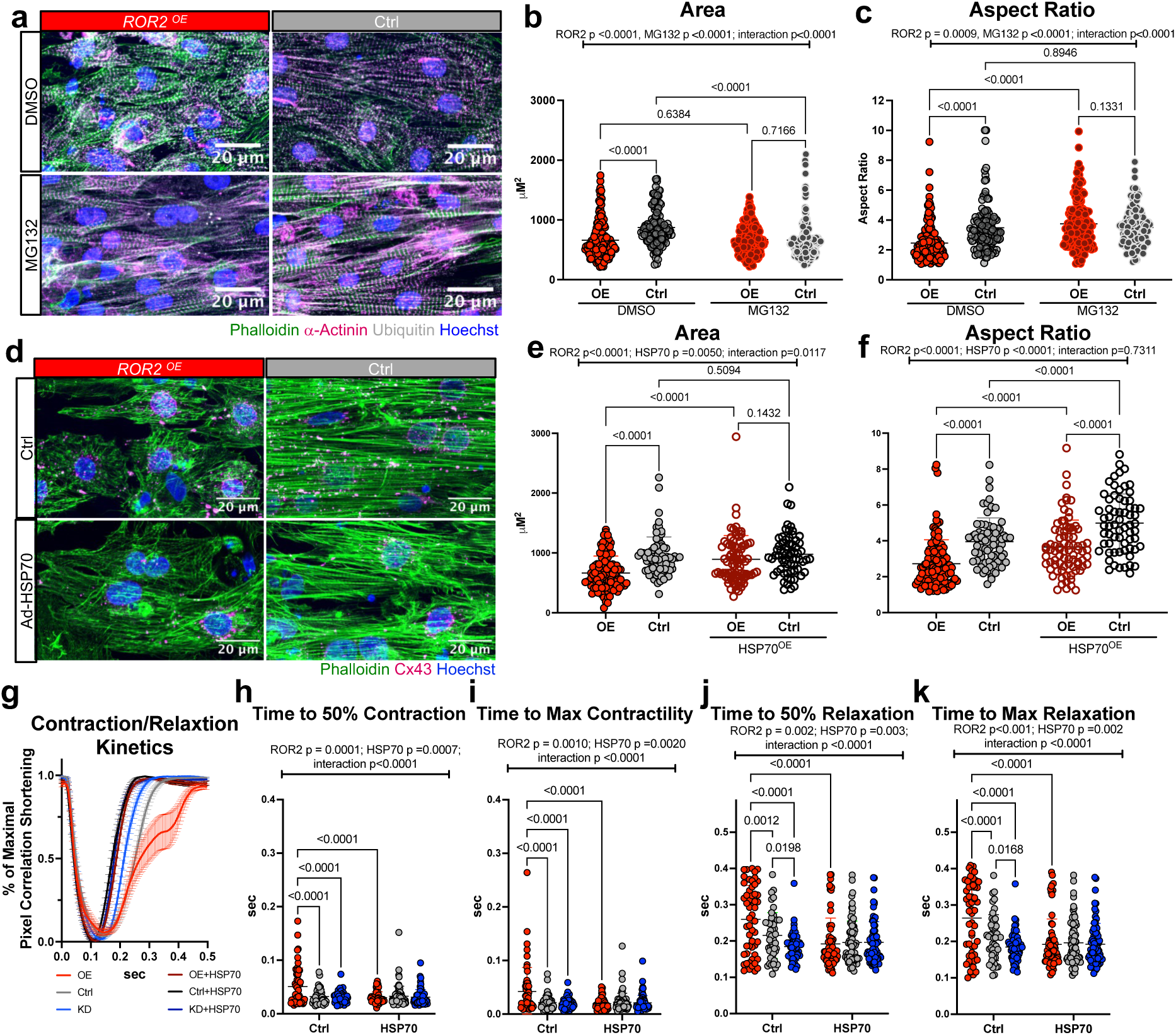
Proteostatic disruption is responsible for ROR2-mediated cardiomyocyte structural and functional defects. The impact of proteasome inhibition (**a-c**; MG132 1μM for 24 hours, n=129-183/group) and increased HSP70 expression (**d-f**; 24 hours of adenoviral expression of *Hspa1b*, n=65-111/group) on ROR2^OE^-mediated change in cell area and aspect ratio was evaluated as described above. Using a Cytocypher pacing apparatus, the impact of ROR2^OE^ and ROR2^KD^ on contractile and relaxation kinetics (**g-k**) were measured using CytoMotion Pixel Correlation with IonOptix software (n=48-75/group from three separate NRVM isolations). Statistics by Two Way ANOVA without correction (**a-f**) or with two-stage step up Benjamini correction (**i-l**).

We next tested the impact of ROR2 modulation on cardiomyocyte contractile properties. Using an IonOptix Cytocypher pacing apparatus, we found that ROR2^OE^ significantly slowed contractile and relaxation kinetics, and that ROR2^KD^ accelerated relaxation (**Fig. 4g-k**).

Strikingly, HSP70^OE^ accelerated contraction and relaxation kinetics in both ROR2^OE^ and control NRVMs to match ROR2^KD^ (**Fig. 4g-k**). Taken together, these data demonstrate that disruptions in proteostatic balance—including UPS-mediated turnover and folding capacity—are primary mechanistic mediators of ROR2-responsive effects on NRVM morphology and function.

### ROR2 regulates the cardiomyocyte proteome

In light of the profound effects of ROR2 on proteostatic balance and evidence that ROR2 impacts protein synthesis and turnover in a selective manner, we next sought to characterize the interaction between ROR2 expression and proteasome inhibition on the global proteome (**Fig. 5a-c**). By principal component analysis plot, we observed tight clustering by groups with PC1 correlating to MG132 and PC2 correlating to ROR2^OE^ effects (**Fig 5c**). Accordingly, ROR2^OE^ dramatically altered the NRVM proteome, which was broadly reversed by proteasome inhibition (**Fig 5a/b**). We considered whether intrinsic protein characteristics contributed to ROR2^OE^-mediated effects on the global NRVM proteome. Using a comprehensive dataset of protein half-lives in mouse fibroblasts and known molecular masses as indicators of stability and complexity, we found that upregulated proteins (adj P<0.05) were smaller and exhibited longer predicted half-lives compared to downregulated proteins (65.8 kDa vs. 87.5 kDa and 134 vs. 55 hours, p<0.0001, both, **Supplementary Fig a/b**).^31^ However, the magnitude of up- or down-regulation did not correlate directly to protein mass or predicted half-lives, suggesting additional levels of regulation **(Supplementary Fig 4c-f**). ROR2^OE^ did not significantly affect the expression of proteasome subunits, consistent with similar PSMB5 levels by western blot as noted above. In contrast, we found ROR2^OE^ significantly altered E3 ubiquitin ligase expression (20 upregulated and 26 downregulated), some of which was reversed by proteasome inhibition, indicating substantial protein substrate-specificity for ROR2-driven UPS-mediated clearance (**Fig**. **5d****/e**, Supplementary Data 20 and 21). These included upregulation of OZZ-E3 (*Neurl2*)—a ubiquitin ligase that targets β-catenin specifically at the intercalated disc—and ITCH—which targets the non-canonical WNT signaling protein DVL (**Fig 5d**). PDZRN3, an E3 ligase that was recently identified as a likely indirect ROR2 phosphorylation target and which plays a role in postnatal sarcomerogenesis and ICD formation, was also significantly reduced.^32,33^ In contrast, we observed no changes in the components of the E3 ligase complex SCF^β-TRCP^ that targets the transcriptional pool of β-catenin, which is consistent with no substantial changes in canonical WNT signaling by RNAseq (**Supplementary Fig. 4a**).

**Fig. 5.**
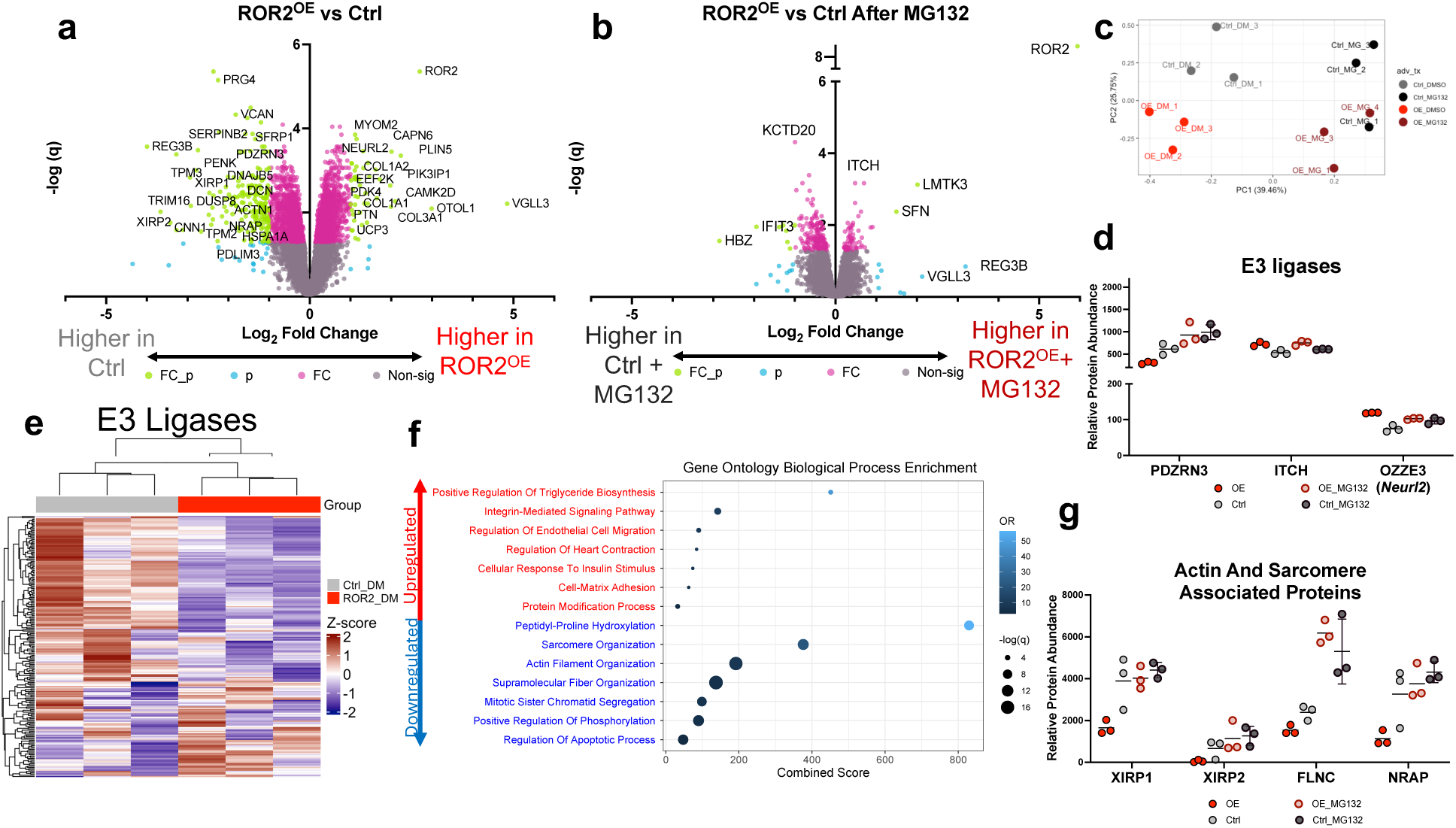
ROR2 mediated proteostatic disruption impacts cardiomyocyte structural proteins. The interaction of ROR2^OE^ and proteasome inhibition on the impact of the NRVM proteome was investigated by TMT-labeled mass spectrometry-based proteomics using Voom/Limma to evaluate differential protein abundance, with visualization by volcano (**a/b**) and PCA plots (**c**). Normalized abundance of representative (**d**) E3 ligases is displayed by dotplot and heatmap (**e**) using a previously published comprehensive list of human E3 ligases.^66^ (**d/e**) Differential protein abundance enrichment analysis was performed for GO Biological Processes (**f**). Representative change in actin and sarcomere associated proteins with ROR2^OE^ and MG132 are displayed by dotplot (**g**).

Pathway enrichment analysis revealed that proteins which were upregulated by ROR2^OE^ were enriched for multiple biological processes including triglyceride biosynthesis, extracellular matrix proteins, integrin-mediated signaling, heart contraction, and response to insulin. Notable upregulated proteins included myogenic differentiation factors VGLL3 and MEF2A—consistent with an immature state—and regulators of calcium homeostasis PLN, CAMK2D, and CAMK2A—which may contribute to altered contractile kinetics. Moreover, in the setting of junctional proteins being mislocalized in ROR2^OE^ by microscopy, the increase in integrin associated proteins may reflect a compensatory attempt to establish cell-cell junctions.

Interestingly, of 426 proteins with at least logFC >0.5 upregulation (adj p<0.05), 60% were significantly decreased by proteasome inhibition compared to 2% that were further increased. Downregulated proteins were enriched for regulators of proline hydroxylation, sarcomere and actin filament organization, supramolecular organization, and inhibitors of apoptosis (**Fig. 5f**). Mirroring the behavior of upregulated proteins, 60% of the 559 downregulated proteins (logFC < -0.5, adj p<0.05) were significantly increased by MG132 with 6% demonstrating further reduction. Importantly, reversal of downregulated proteins included many actin cytoskeleton and sarcomere associated proteins including: TPM1/2/3, XIRP1/2, MYOZ2, LMOD2, FLNC, NRAP, and TNNT2 (**Fig. 5g** and **Supplementary Fig. h/i**). Thus, we conclude that ROR2-mediated disruption of the cardiomyocyte proteome impinges on cardiomyocyte differentiation, structural, functional, and adhesion proteins, that are largely dependent on altered UPS activity.

### Cardiac ROR2 expression causes RV failure and proteostatic disruption

Given the marked effects of ROR2^OE^ in cardiomyocytes in cell culture, we next sought to test the effect of ROR2 expression in the RV *in vivo*. To this end, we generated cardiac ROR2^OE^ mice using AAV9 to deliver HA-tagged ROR2 versus GFP in 4-week-old male and female mice (**Fig. 5a**). At 8 weeks after injection, we observed robust RV and LV ROR2 expression, which localized to the cardiomyocyte membrane and cytoplasm (**Fig. 6b, Supplementary Figs. 1a and 5a**). ROR2^OE^ led to significant RV hypertrophy in male mice, marked by an increase in RV mass, RV wall area, and RV cardiomyocyte cross-sectional area by minimum Feret Diameter (**Fig. 6b-h**). Echocardiography (n=8/group, 50% male) revealed biventricular dilation and systolic dysfunction in male ROR2^OE^ mice (**Fig. 6i-l** and **Supplementary Table 3**). Cardiac catheterization revealed a male-specific increase in RV end diastolic pressure (**Fig. 6m**).

**Fig. 6.**
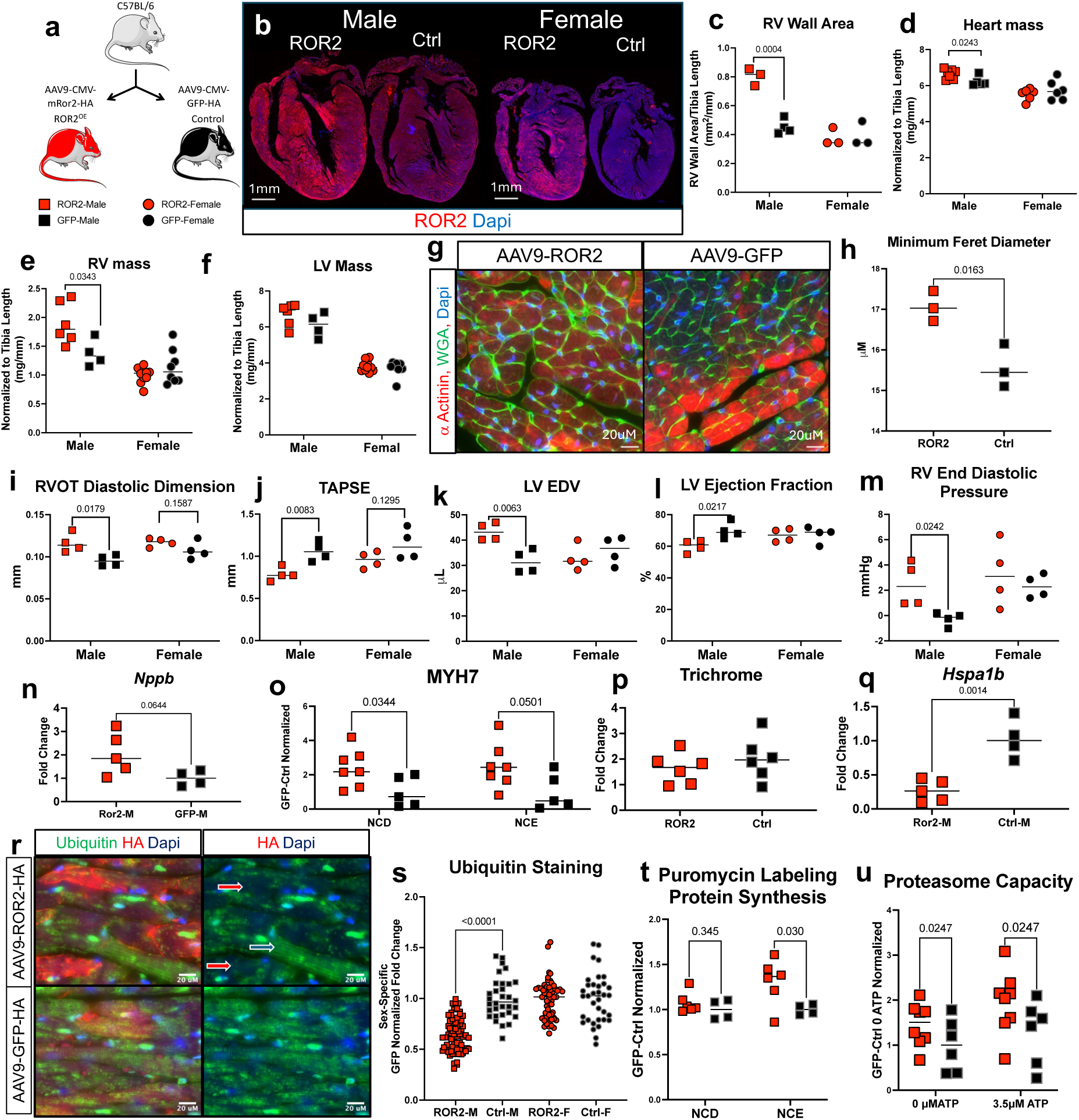
Cardiac ROR2 induction is sufficient to cause RVF and proteostatic disruption in mice. (**a**) The impact of cardiac ROR2^OE^ was tested using AAV9 delivery of HA-tagged *Ror2* vs GFP in male and female mice at age 4 weeks with phenotyping performed at age 12 weeks. Cardiac ROR2^OE^ caused a male predominant phenotype with increase in RV wall area **(b** representative 10X magnification and **c**) quantified using FIJI with color deconvolution to identify epi and endocardial borders. Total heart mass after trimming great vessels and RV free wall and LV mass were measure after removal of atria and separation of RV free wall (**d-f**). RV cardiomyocyte cross sectional area was measured using HeartJ^67^ FIJI package on WGA stained section and quantifying the minimum Feret Diameter (**g/h**, 40X magnification). Echocardiography was performed to assess biventricular size and systolic function (**i-l**) and jugular catheterization was performed to assess RV end diastolic pressure (**m**). Expression of *Nppb* was measured by qPCR (**n**) and fetal MYH7 was measured by western blot (**o**). Fibrosis was measured using trichrome staining of RV free wall tissue (**p**). Proteostatic characterization including assessment of *Hspa1b* by qPCR (**q**), ubiquitin immunohistochemistry of HA-tag (i.e. AAV infected cardiomyocytes) (**r/s**, 60X magnification), puromycin labeling to measure protein synthesis rates using a 1-hour puromycin pulses by intraperitoneal injection (**t**) and proteasome capacity using Suc-LLVY-AMC (**u**). Statistical comparisons performed using t-test or Mann Whitney depending on normality, (n=3-8 mice/group and 32-72 RV cardiomyocytes/group for panel **s**).

ROR2^OE^ male mice also exhibited induction of *Nppb* mRNA and fetal MYH7 **(Fig. 6n/o**) without causing an increase in fibrosis (**Fig. 6p**). The cardiac dysfunction seen in male ROR2^OE^ mice was accompanied by proteostatic disruptions similar to those seen in NRVMs including lower *Hspa1b* mRNA, decreased cardiomyocyte ubiquitin staining, increased protein synthesis, and increased proteasome capacity **(Fig. 6q-u** and **Supplementary Table 4**). We also observed reduced peripheral β-catenin in RV cardiomyocytes consistent with disrupted cell-cell junctions that mirror effects seen in NRVMs (**Supplementary Fig. 5b/c**). We conclude that ROR2 expression is sufficient to drive RV failure *in vivo* in male mice, concomitant with disruption of proteostasis, whereas female ROR2^OE^ mice were generally much less affected.

### Suppression of ROR2 induction in RV pressure overload abrogates RV failure and proteostatic disruption

The results above demonstrated that ROR2 expression was sufficient to cause RV dysfunction and disrupt proteostasis. We next sought to test if ROR2 expression was necessary for RVF in the PAB mouse model and its role in proteostasis. We first evaluated the proteostatic signature of PAB-induced RVF, noting that PAB increased RV protein synthesis, HSP70, and total ubiquitin—with K48 ubiquitin and PSMB5 increased in the NCE fraction—and proteasome capacity (**Fig. 7a-g** and **Supplementary Table 5**). Thus, the PAB RVF mouse model demonstrates clear proteostatic disruption with an increase in both protein synthesis and UPS-mediated turnover, to some extent analogous to that found with ROR2^OE^. Next, to determine if ROR2 induction in PAB contributes to RVF and proteostatic disruption, we injected male mice with AAV9 to deliver *shRor2* (PAB^shRor2^, n = 15) or scrambled shRNA (PAB^shScr^, n=15) prior to PAB. Since ROR2 localization in the NCE fraction correlated to PAB severity, we assessed the relative ROR2 knockdown in NCD and NCE fractions and found PAB^shRor2^ had ∼50% lower NCE and ∼25% lower NCD ROR2 in the RV (**Fig. 7i/j**). Strikingly, PAB^shRor2^ mice demonstrated less severe RV dilation, decreased incidence of tricuspid insufficiency, better RV systolic function measured by TAPSE (**Fig. 7j-l**), improved diastolic function by e’ (**Fig. 7m**), less RV fibrosis, and reduction in RV wall area by histology (**Fig. 7k-p** and **Supplementary Table 6**). ROR2 knockdown also ameliorated the proteostatic dysfunction in PAB with decreases in protein synthesis, HSP70, and K48-linked ubiquitin without affecting proteasome capacity (**Fig. 7q-t**), partially reversing the PAB-induced defects. Taken together, these data demonstrate that ROR2 knockdown protects from pathologic RV remodeling and improves RV function in response to pressure overload, concomitant with partial reversion of altered proteostasis.

**Fig. 7.**
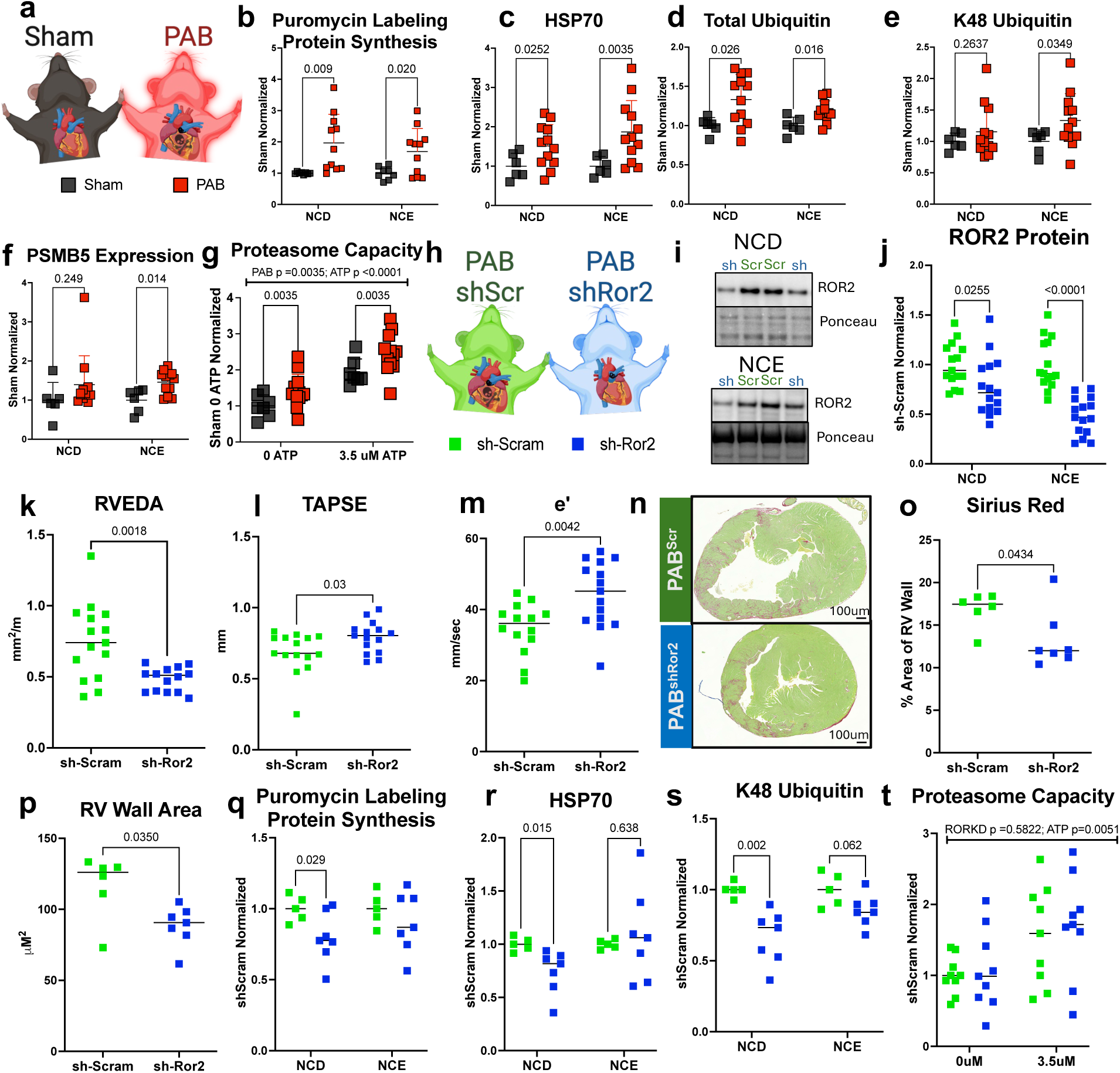
ROR2 knockdown improves RVF and proteostatic burden in RV pressure overload. Proteostatic characterizations were performed for PAB and sham mice (**a**) by puromycin labeling to measure protein synthesis (**b**), expression of HSP70 (**c**), total ubiquitin (**d**), K48-linked ubiquitin (**e**), and PSMB5 expression (**f**) by western blot in NCD and NCE fractions. Proteasome capacity was measured using Suc LLVY-AMC (**g**). ROR2^KD^ was achieved by AAV9-delivery of *Ror2* targeting shRNA (PAB^shRor2^) delivered by retroorbital injection 4-weeks prior to PAB/sham, which reduced NCD (∼25%) and NCE (∼50%) ROR2 expression compared to scrambled shRNA (PAB^Scr^; **h-j**). Echocardiography was performed 2 weeks after PAB to assess RV remodeling, systolic and diastolic function (**k-m**). Histology using Sirius red of RV free wall and FIJI to assess RV area and fibrosis percent using colour deconvolution (**n-p**). Proteostatic state in PAB^shScr^ compared to PAB^shRor2^ was characterized using puromycin labeling, K48 ubiquitin and HSP70 expression, and proteasome capacity by Suc-LLVY-AMC as above (**q-t**). Statistics by t-test or Mann Whitney depending on normality (n=6-7 for Sham vs PAB n=9-12 and n=5-15 PAB^shScr^ vs n=7-14 for PAB^shRor2^), or Two-Way ANOVA for proteasome capacity.

### ROR2 expression in human RVF correlates to proteasome capacity

Finally, we sought to determine in human RVF the relationship between ROR2 and proteostasis. To do so, we leveraged a collection of RV tissue obtained from non-failing organ donors compared to DCM RVs collected at transplant. DCM RVs were stratified by having preserved RV function versus RVF, and RVF hearts were further stratified by the level of NCD and NCE ROR2. We evaluated for demographic, clinical, and anthropometric balance across all relevant comparisons (**Fig 8a** and **Supplementary Tables 7-10**). We observed that high ROR2 expressing RVF, stratified by NCE or NCD fractions, demonstrated significantly increased proteasome capacity at 0, 3.5, and 7 μM ATP (**Fig 8, Supplementary Fig 6a**, and **Supplementary Table 11**; p<0.05). Interestingly, NCE ROR2 showed significant Pearson correlation to proteasome capacity at each of the tested ATP levels (r=0.68 – 0.82, p≤ 0.0008; **Fig 8c** and **Supplementary Tables 12**), whereas NCD ROR2 showed no clear relationship. We also assessed the abundance of other proteostatic markers and found that high NCE ROR2 associated with a trend for decreased NCE HSP70 and that high NCD ROR2 associated with increased NCD K48 ubiquitin (**Supplemental Fig 6b-e**). However, we found no clear patterns for PSMB5 and total ubiquitin (**Supplemental Figure 6f-i)**. We conclude that in human cardiomyopathy, as in mice, high ROR2 expression—particularly NCE ROR2—is associated with an increase in proteasome capacity.

**Fig. 8.**
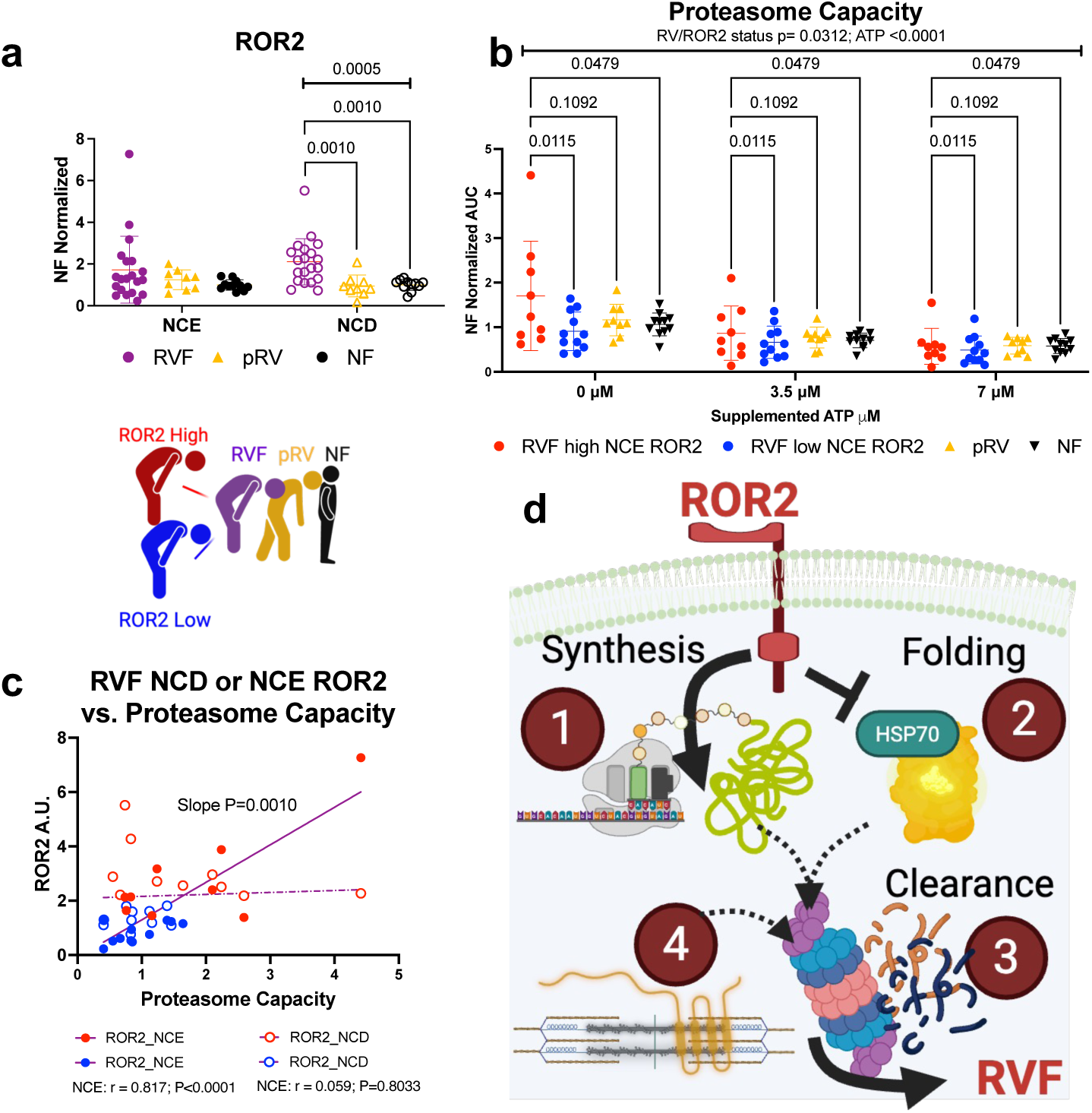
ROR2 expression is associated with proteostatic disruption in human RVF. (**a**) ROR2 expression in NCE (filled in symbols) and NCD (open symbols) fractions was measured in the RVs from humans with RVF from dilated cardiomyopathy with (purple), preserved RV function (orange triangles, pRV), or nonfailing (NF, black). RVF samples were further stratified by high (red symbols) and low (blue symbols) NCE/NCD ROR2. (**b**) Proteasome capacity was compared across human RV functional groups. (**c**) Within RVF, NCD/NCE ROR2 levels were assessed for Pearson correlation against proteasome capacity (noted below graph) and for differences in linear slopes. Statistics by Kruskal-Wallis (**a**) or Two Way ANOVA (**b**) with two-stage step up Benjamini correction (n=9-11/group). (**d**) Taken together, our work reveals a model wherein high ROR2 expression in cardiomyocytes disrupts proteostasis with increased protein synthesis (1) and reduced folding (2) that results in a poor-quality proteome, leading cardiomyocytes to clear proteins by the UPS (3), which disrupts cardiomyocyte structure and function (4).

## Discussion

RVF remains a widespread, challenging problem for which there are no proven therapies.^14^ The breadth of LVF medications have not shown clear benefit for RVF, indicating the presence of distinct RVF mechanisms.^7–13,34^ In severe human RVF, we found that the developmentally restricted noncanonical WNT receptor ROR2 is pathologically reactivated by both cardiomyocytes and fibroblasts.^16^ We now find this expression pattern is recapitulated in mouse pressure overload RVF, which contrasts with a prior report in mouse LVF for which induced ROR2 is restricted to noncardiomyocytes.^29^ Using NRVMs, in vivo murine models, and human heart biospecimens, we now reveal a previously unrecognized role for ROR2 regulating cardiomyocyte proteostasis with implications on contractile function and structure. Interestingly, in mouse and human RVF, NCE localizing ROR2 demonstrates a stronger association than NCD ROR2 with pathologic RV remodeling and proteostatic disruption. Our data point to a model wherein ROR2 promotes protein synthesis without sufficient folding capacity (**Fig. 8d**).

Although UPS-mediated clearance is upregulated in this setting, this is either not sufficient to meet the demands or, potentially, chronically maladaptive and ultimately converges on cardiomyocyte functional compromise. Thus, we propose that addressing ROR2-mediated proteostatic disruption represents a potential novel therapeutic avenue for RVF, for which no proven therapies currently exist.

ROR2 is a noncanonical cell surface WNT receptor pseudokinase that signals through at least three broad pathways: RAC/RHO regulation of cell polarity, activation of a JNK/cJUN phosphorylation cascade, and activation of Ca^2+^-responsive targets (e.g. PKC, CAMKII).^35^ The molecular and functional impact of pathogenic ROR2 has been best studied in cancer, for which ROR2 causes context-specific pleiotropic phenotypes with mixed impact on cancer cell death, invasion, and metastasis resulting in both positive and negative associations with patient mortality.^36,37^ Little is known about the role of ROR2 in cardiac pathology, but upregulation of ROR2 leading to proteostatic disruption in RVF fits within the well-recognized paradigm of fetal gene program reactivation during heart failure.^38,39^ As the first functional organ, proliferating fetal cardiomyocytes must maintain a high quality pool of contractile proteins, including some of the largest in the proteome (e.g. Titin, Obscurin, Dystrophin, etc.), which is energetically demanding and requires high protein synthesis matched to a robust folding capacity.^40–42^ During human heart development, *ROR2* and *HSPA1A/B* (HSP70) mRNA levels are positively correlated and taper during later prenatal stages, consistent with amplified but balanced proteostasis of the immature heart.^43^ Postnatally, in mice we found age-dependent reductions in ventricular HSP70 and ROR2, with reductions in ROR2 preceding HSP70. In mouse PAB RV, translation, folding, and proteasome capacity are all increased, but the presence of ubiquitin accumulation suggests there is imbalance in these three components that favors excess translation.^42^ Moreover, given that ROR2 knockdown in PAB improves RV remodeling and function while most substantially affecting protein synthesis, attenuating the translational hyperactivation of RVF may be the critical factor to restore proteostatic balance. Prior work has shown that slowing translation with rapamycin can improve translational fidelity in mTOR activated embryonic fibroblasts and can rescue multiple preclinical LVF models.^44–46^ In kind, rapamycin was shown to abrogate PAB-induced RV hypertrophy and contractile dysfunction in rats.^44^ Although protein translation and UPS capacity were not measured, rapamycin did further increase PAB-induced ubiquitin accumulation suggesting a decrease in UPS-mediated clearance, which may be the result of slowing translation leading to improved overall proteome quality.

Proteostatic disruption is strongly implicated in heart failure, and its role in RVF is increasingly recognized, yet developing targeted therapies has remained elusive.^47–49^ Restoring proteostatic balance would require precise control of synthesis, folding, and turnover, or a more practical strategy of eliminating a pathogenic upstream regulator. Before this study, few links connected ROR2—or noncanonical WNT signaling more broadly—to proteostasis. In cultured ovarian carcinoma cells that typically express low ROR2, adenoviral ROR2 delivery activated the unfolded protein response and triggered apoptosis.^50^ These ROR2 effects in ovarian carcinoma were rescued by blocking induction of the endoplasmic reticulum stress protein IRE1a. In NRVMs, treatment with WNT5A—the only known ligand for ROR2—was shown to promote protein synthesis and drive cardiomyocyte hypertrophy.^51,52^ These WNT5A effects could be blocked by preventing the membrane localization of the ROR2 binding partner VANGL2 or by inhibiting JNK. Although JNK/cJUN are not classic regulators of protein synthesis, extensive evidence shows crosstalk with mTOR and ERK. For example, both canonical and noncanonical WNT activation can inhibit the signaling hub protein GSK3 through phosphorylation.^53,54^ Inhibiting GSK3 prevents it from phosphorylating several key substrates including β-catenin—(leading to canonical WNT activation), TSC2 (activating mTORC1), and Raptor (suppressing mTORC1).^55–57^ JNK can also activate mTORC1 through Raptor phosphorylation, as observed in HEK293 cells under osmotic stress.^58^ ERK signaling provides an additional link. In A375 melanoma cells which have low endogenous ROR2, induction of ROR2 caused ERK hyperactivation and cell death.^36^ In isolated mouse cardiomyocytes, ERK1/2 activation and HSP70 expression show reciprocal antagonism, and HSP70 knockdown was sufficient to cause contractile and relaxation deficits similar to those we observed with ROR2^OE^ NRVMs.^59^ Thus, ROR2 could lead to a pathologic increase in protein translation, thereby leading to proteostatic disruption in cardiomyocytes, by activating mTOR or ERK signaling through modulating activity of other true kinases (e.g. JNK/cJUN or GSK3)

Using orthogonal methods, we found that ROR2 can localize to the nucleus in the immature heart and RVF. Nuclear enriched ROR2 in the NCE fraction strongly correlated to pathogenic RV remodeling and proteostatic disruption. The functional role of nuclear ROR2 remains poorly characterized, but in cancer nuclear ROR2 correlates to worse disease with increased metastasis in melanoma and invasiveness in pancreatic cancer.^26,28^ These pathophysiologic associations with nuclear ROR2 underscore the importance of understanding the regulatory mechanisms and functional roles (e.g. transcriptional modulation, scaffolding, etc.) of nuclear localizing ROR2, which may further enable selective therapeutic targeting. For example, ERK1/2 homeostatic and pathogenic roles are compartmentalized into the cytoplasm and nucleus, respectively, and blocking its nuclear localization was shown to prevent pressure induced cardiomyocyte hypertrophy and heart failure.^60,61^

### Limitations

More work will be necessary to establish the therapeutic potential for targeting ROR2 in RVF. While we have not observed sexual dimorphism for the impact of ROR2 in humans, we found that overexpressing ROR2 in female mice led to a milder phenotype compared to males. This sex difference in mice may fit with prior reports showing female mice are protected from proteostatic disruption through higher basal chaperone expression and UPS capacity.^62–64^ We focused our in vitro mechanistic studies in cardiomyocytes, but ROR2 upregulation was also observed in fibroblasts of human and mouse RVF. Importantly, in a prior mouse model of left ventricular pressure overload where ROR2 induction was restricted to fibroblasts and leukocytes, inducible total body combined ROR1/ROR2 knockout was lethal, diminished early myofibroblast activation, and increased the abundance of inflammatory cells in the LV.^29^ The differences between that study and ours may be due to multiple factors including ubiquitous knockout of both genes compared to knockdown of only ROR2 and by differences in the models (RV vs LV pressure overload and the implicated cell populations) that may alter the impact of loss of ROR2. Since ROR2 is known to be pleiotropic, understanding the role of ROR2 across the spectrum of fibroblast activation states could have significant implications for its therapeutic targetability.^29, 65^ Finally, our analysis on the relationship between ROR2, RVF, and proteostasis in humans reflects a single, uncontrolled timepoint which prevents us from measuring dynamic processes (e.g. UPS flux, protein synthesis rates) that would more fully capture this relationship.

### Conclusions

In summary, we find compelling evidence to implicate ROR2 in RVF pathogenesis. Pathologic reactivation of ROR2 causes proteostatic disruption in the RV, leading to RVF. In humans, ROR2 is otherwise broadly absent from healthy mature tissues with its only other well recognized induction occurring in cancer.^23^ Thus, we conclude that targeting ROR2 offers a potential therapeutic strategy to restore proteostatic balance and treat RVF, for which no specific therapies currently exist.

## Acknowledgements

We thank Julian Mintseris, Rachel Rodrigues, and Jonathan Van Vraken at Thermo Fisher Scientific Center for Multiplexed Proteomics at Harvard Medical School for support in performing proteomics. We thank the Day Lab at University of Pennsylvania for kindly providing adenovirus to express GFPμ. The mouse PAB and echocardiography were performed by the Rodent Cardiovascular Phenotyping Core (RRID: SCR_022419) at the University of Pennsylvania supported by the Penn Cardiovascular Institute and NIH S10OD016393. Figures were generated with assistance of BioRender.

## Author Contributions

H.H., J.J.E., and Z.A. developed the concept and designed the experiments. J.J.E, Z.A., and B.L.P. supervised and funded the project. J.J.E., H.H., K.U., E.A.S., G.U., S.K., and S.W. conducted NRVM experiments. J.J.E., H.H., D.E., and R.T.S. conducted NRVM contractility/relaxation experiments. J.J.E and H.H. performed mouse experiments. J.J.E, T.D., and C.S.C. conducted luciferase refolding and ^35^S-Met labeling assays. J.B. and Y.Y. performed RNAseq analysis. L. Lai, J.G., and S.S. performed mouse surgeries and physiologic characterization. K.C.B. and K.M. supported human heart tissue procurement and clinical characterization. H.R. contributed to microscopy image analysis. L.Li performed tissue histology. All authors contributed to the manuscript preparation and approved the final version of the manuscript.

## Corresponding author

Correspondence to Jonathan J. Edwards and Zoltan Arany.

