## Supplemental figures and tables for "ROR2 drives right ventricular heart failure via disruption of proteostasis"

**Manuscript Supplement:**

I. Detailed Methods

II. Supplementary Figures and Legends

III. Supplementary Tables

IV. List of Source Data (see separate Excel file)

### **Detailed Methods**

#### In vitro experiments

*NRVM culture*

All animal work was performed under University of Pennsylvania Institutional Animal Care and Use Committee approved protocols (#805255 and #805309). Neonatal rat ventricular myocytes (NRVMs) were isolated and maintained in culture as previously described.[21] Briefly, 1-2 day old Sprague Dawley rat pups were anesthetized on ice, rapidly decapitated and their hearts were excised and placed in chilled Hanks' Balanced Salt Solution (HBSS, Sigma-Aldrich). Ventricular tissues were pooled from a single litter, minced and digested in HBSS with trypsin (Worthington Biochemical) and benzoase (Sigma-Aldrich). Cells were centrifuged at 500g x15 minutes to remove residual trypsin and resuspended in serum-containing NRVM media (DMEM ThermoFisher, 5% FBS, 12.5 mM HEPES (UPenn Cell Center), 4mM Aln-Gln (Sigma-Aldrich), 0.1 mg/mL primocin. Cells were preplated for 2h in cell-culture treated flasks to remove the majority of the cardiac fibroblasts. NRVMs were then plated on plasma treated (PlasmaFlo, Harrick Plasma), collagen-coated nano-patterned culture dishes or coverslips (Curi Bio) at 150,000 cells/cm$^2$. After overnight attachment in serum-containing media, the cells were serum starved for 48h before further treatment (serum-free media: DMEM, 1X insulin-transferrin-selenium (ThermoFisher), 12.5 mM HEPES, 4mM Aln-Gln, 0.1 mg/mL primocin, 0.3% BSA. NRVMs were maintained in serum-free NRVM media throughout remaining experiments with media changes every 48 hours. Where indicated, NRVMs were treated with MG132 (1uM or 20μM).

*Generating ROR2-modified NRVMs*

ROR2 overexpression ($ROR2^{OE}$) NRVMs were generated using adenovirus expressing human *ROR2* under a CMV promoter with an eGFP reporter under a bicistronic CMV promoter (Vector Biolabs ADV-221490) at a multiplicity of infection (MOI) of 100. $ROR2^{KD}$ NRVMs were similarly generated using U6 promoter to express *Ror2* targeting shRNA with a CMV-eGFP (shADV-252440) reporter and control NRVM received virus expressing eGFP alone (cat 1060). For cell morphology evaluation after MG132 and for proteomics analysis, $ROR2^{OE}$ (cat ADV-221490) and control (cat 1300). NRVMs were collected for further analysis 72-hours after adenovirus infection. For experiments with proteasome inhibition, NRVMs were treated with 20µM MG132 in DMSO for short exposures or 1µM DMSO for 24 hours prior to collection. For experiments with concomitant HSP70 overexpression, 48 hours after initial adenovirus treatment NRVMs were then incubated with Ad-mCherry-h-*HSPA1B* (Vector biolabs: ADV-232159) at MOI of 100 for 24 hours prior to cell collection.

shRNA *Ror2*:

5'-CCGG-CGTGGTGCTTTACGCAGAATACTCGAGTATTCTGCGTAAAGCACCACG-TTTTT-3'

shRNA scramble:

5’-GACACGCGACTTGTACCACTTCAAGAGAGTGGTACAAGTCGCGTGTCTTTTTTACGCGT-3’

*NRVM RNAseq*

NRVMs lysates were collected in RNAlater$^{TM}$ (ThermoFisher) stabilization fluid and total RNA were extracted using RNeasy Mini Kit (Qiagen). RNA samples were diluted to 100 ng/µL by DEPC water using Qubit fluorometer (Thermo Fisher Scientific Waltham, MA). RNAseq was performed by Azenta with PolyA selection on an Illumina HiSeq using 150 bp paired end reads. The paired-ended RNAseq reads were aligned to the rat reference genome (Rnor-6.0) using STAR (version 2.7.9a)(https://pubmed.ncbi.nlm.nih.gov/23104886/). We used Fastqc (version 0.11.7) (https://www.bioinformatics.babraham.ac.uk/projects/fastqc/) to check the quality of the reads, Picard (https://broadinstitute.github.io/picard/) to mark the duplicates, compute the RNAseq metrics and estimate the library complexity. We then calculated the raw read counts using the Rsubread package (version 2.0.3) (https://academic.oup.com/nar/article/47/8/e47/5345150). Genes with 25% of samples having a CPM <1 were considered as low-expressed and were removed from further analysis. We used the voom function from the Limma package (version 3.56.1) to transform and normalize the data, and the toptable function to output the differential analysis result.

Gene set enrichment analysis was performed using Enrichr with genes exhibiting an adjusted $p<0.05$ comparing ROR2$^{OE}$ and ROR2$^{KD}$. To specifically compare these differential expression data to the cardiac-specific canonical Wnt targets, we downloaded the RNAseq counts file from the GEO Dataset GSE121234 which includes data from a cardiomyocyte-specific non-degradable β-catenin gain of function mouse (Myh6$^{Mer\text{-}Cre\text{-}Mer}$x β-catenin$^{\Delta Ex3}$) and wild type P6 ventricular myocardium. We performed a similar Limma/Voom differential expression analysis and filtered for all commonly identified genes as the background list and adj $p<0.1$ for statistically significant genes.

*NRVM Immunofluorescence Microscopy*

NRVMs were washed in PBS and fixed in 4% PFA (Electron Microscopy Sciences) for 15 minutes, washed twice with PBS, permeabilized in 0.1% TritonX-100 for 10 minutes at RT, and washed twice with PBS. NRVMs were blocked using Seablock (Abcam) for 1 hour at RT and then incubated in primary antibody diluted in Seablock for 48 hours at 4°C. NRVMs were washed twice with PBS and then incubated in secondary antibody for 24 hours at 4°C. NRVMs were washed twice in PBS, incubated in 1:1000 Hoechst 33342, trihydrochloride, trihydrate (Invitrogen H3570) for 10 minutes at RT, washed twice with PBS, and then mounted onto coverslips using Prolonged Diamond.

NRVMs were imaged on a Zeiss 880 Airyscan confocal microscope operating on an Axiovert Z1 inverted microscope equipped with Plan-Apochromat x63 oil 1.4 numerical aperture objective. Image analysis was performed using ZEN Black software for Airyscan processing to integrate 32 separate sub-resolution detectors in the Airyscan detector and subsequent deconvolution. A z-stack of four 1μM thick slices and 2x2 tiles were captured for each coverslip acquired at regions with representative cell density. Stitching was performed using ZEN Black software.

Further image processing and quantification was performed using FIJI. Each z-stack was converted to a maximum intensity projection. Every NRVM with complete edges visualized in the image were quantified without exclusion. The alpha-actinin or phalloidin channel was used to trace the cell border to generate regions of interest (ROIs). The peripheral ROI for each cell was generated using a FIJI macro to reduce the cell ROI by 2μM in all directions (“AutoShrink”, enlarge=-2). NRVM structure was quantified using area, major and minor axes, and aspect ratio. Staining was measured using mean intensity for the entire ROI. To assess peripheral staining enrichment, the difference in raw integrated density between the full ROI and inner ROI was normalized to the difference in area between these ROIs and by the mean intensity for the inner ROI.

*NRVM Protein Extraction*

NRVMs were washed in PBS and total protein was extracted by scraping and pipetting in extraction buffer: 8M urea, 2M thiourea, 3% sodium dodecylsulfate, 75 mM dithiothreitol, and 50 mM Tris pH 7.5.[68] The extraction buffer was deionized using mixed-bed resin (Biorad, catalog no 1426425) and supplemented with protease/phosphatase inhibitor (Halt™ Protease and Phosphatase Inhibitor Cocktail, ThermoFisher). Cell lysates were mixed 1:1 with 50% glycerol in water.

*Protein synthesis rate quantification*

NRVM translation rate was measured by labeling with the tyrosyl-tRNA analogue puromycin labeling.[21] After generating ROR2$^{OE}$, GFP-ctrl, and ROR2$^{KD}$ NRVMs with or without concomitant HSP70$^{OE}$, cells were incubated with low dose (1 μM) puromycin for 1 hour at 37C. As a negative control, some wells were simultaneously incubated with cycloheximide (20 ug/mL). Cells were washed with PBS containing cycloheximide and lysates were collected using thiourea. Puromycylated proteins were quantified by western blot using antipuromycin antibody and normalized to total protein loading by ponceau. Protein translation was also measured similarly using radiolabeled methionine ($^{35}$S-Met) for 1 hour.[69] $^{35}$S-Met labeled proteins were quantified using autoradiography and normalized to ponceau.

*Proteasome activity quantification*

The ubiquitin proteasome system (UPS) in NRVMs was evaluated in two ways: by fluorogenic substrate Suc-LLVY-AMC and with puromycin pulse chase. [70]

*Chymotrypsin-like proteasome capacity*

Crude lysates from ROR2$^{OE,}$ ROR2$^{KD}$, and GFP-control NRVMs were collected in NP-40 cell lysis buffer (Invitrogen FNN0021) using sonication at 4$^{o}$C. Cell lysates were clarified by centrifugation at 10,000G x 10 min at 4$^{o}$C, and supernatant was collected. Protein quantification was performed by BCA assay and samples were diluted to 0.8 µg/µL. Chymostrypin-like proteasome activity was directly measured using the fluorogenic peptide substrate Suc-LLVY-AMC as follows: 200µL of assay buffer (50mM HEPES pH7.5, 20 mM KCl, 5mM MgCl2, and 1 mM DTT in ddH20) was added to each well of a 96-well black bottom polystyrene plate (Thermo Scientific, cat 137101). A subset of wells received additional assay buffer or supplemented ATP (final concentration of 3.5 or  7 µM), and some wells received the chymotrypsin-like specific inhibitor Lactacystin (final concentration 18µM). After this, 10µL of sample was added followed by 10 µL of Suc-LLVY-AMC (final concentration 19.5 µM). Technical replicates were performed in duplicate and averaged. Fluorescence was measured every 1 minute for 45 minutes on a BioTek Synergy Neo2 Plate Reader at 37$^{o}$C. The difference in fluorescence intensity without and with lactacystin for each timepoint was plotted, and the area on the under the curve was used to determine proteasome capacity.

Proteasome capacity was measured similarly from mouse and human myocardium with a few modifications. Protein extraction from snap-frozen RV myocardium was performed directly in assay buffer and diluted to 2 µg/µL using BCA assay. Fluorescence was measured every 1 minute for 60 minutes. The area under the curve normalized to control samples for the experiment.

*Puromycin pulse chase*

Puromycin labeling was performed as above, but after the 1-hour labeling wells were washed with PBS and then media was replaced with cycloheximide (20 µg/mL) and either DMSO or MG132 to block proteasome activity (20 µM) for 1 hour at 37°C prior to protein extraction.

*Luciferase refolding assay*

NRVMs cultured on 35-mm nanopatterned dishes NRVM were extracted using 350µL of Passive Lysis buffer (Promega, Inc.) supplemented with 1X protease inhibitor (Halt Protease Inhibitor), and lysates were maintained at on ice.[71] QuantiLum® Recombinant Luciferase protein was 1 mg (supplied as 10 mg/mL) was dissolved in 1mL of passive lysis buffer and heat shocked at 42°C for 15 minutes to denature the protein. The denatured luciferase protein was spiked into the NRVM lysates at a final concentration of 200 ug/mL and incubated at 37°C for 1 hour. 80µL lysate/luciferase renaturing sample was added to a white bottom 96-well plate, then 20µL of luciferase assay reagent was added and mixed. The plate was immediately read on a BioTek Synergy Neo2 Plate Reader at 37°C

*NRVM Contractility and Relaxation Measurements*

NRVM were plated on collagen coated 35mm MatTek glass bottom cell culture dishes for $ROR2^{OE}$, $ROR2^{KD}$, and GFP control with and without HSP70 overexpression as above. NRVMs were paced with the Cytocypher apparatus (IonOptix) using 20 V at 2 Hz. For each replicate, 10-20 cardiomyocytes across multiple regions of each dish were analyzed using CytoMotion Pixel Correlation analysis software without excluding any recording. Every NRVM contractility and relaxation were quantitatively assessed using the time to maximal depression and relaxation velocity as well as time to 10%, 50%, and 90% of peak shortening or return to baseline.

*Mass Spectrometry for Proteomics*

To assess how ROR2 expression affects the proteome in an unbiased manner, tandem mass tag (TMT)-labeled total was performed from $ROR2^{OE}$ and negative-control NRVM lysates according to previously published protocol.[72] Total lysates were collected using TMT-labeling compatible lysis buffer (8M Urea, 200 mM EPPS pH8.5, Roche cOmplete protease inhibitor, Roche PhosSTOP inhibitor, 1% SDS in $ddH_2O$). Samples were analyzed using Orbitrap Eclipse mass spectrometer.

*Sample Preparation for Mass Spectrometry*

Samples for protein analysis were prepared essentially as previously described.[72,73] Following lysis, protein precipitation, reduction/alkylation and digestion, peptides were quantified by BCA assay and 150µg of peptide per sample were labeled with TMTPro reagents (Thermo-Fisher) for

2hrs at room temperature. Labeling reactions were quenched with 0.5% hydroxylamine and acidified with TFA. Acidified peptides were combined and desalted by Sep-Pak (Waters). Peptides from the flow-through were further fractionated for full proteome analysis.

*Basic pH reversed-phase separation (BPRP)*

TMT labeled peptides were solubilized in 5% ACN/10 mM ammonium bicarbonate, pH 8.0 and 300 μg of TMT labeled peptides was separated by an Agilent 300 Extend C18 column (3.5 μm particles, 4.6 mm ID and 250 mm in length). An Agilent 1260 binary pump coupled with a photodiode array (PDA) detector (Thermo Scientific) was used to separate the peptides. A 45-minute linear gradient from 10% to 40% acetonitrile in 10 mM ammonium bicarbonate pH 8.0 (flow rate of 0.6 mL/min) separated the peptide mixtures into a total of 96 fractions (36 seconds). A total of 96 Fractions were consolidated into 24 samples, acidified with 20 μL of 10% formic acid and vacuum dried to completion. Each sample was desalted via Stage Tips and re-dissolved in 5% formic acid/ 5% acetonitrile for LC-MS3 analysis.

*Liquid chromatography separation and tandem mass spectrometry*

Proteome data were collected on an Orbitrap Lumos mass spectrometer (ThermoFisher Scientific) coupled to a Proxeon EASY-nLC 1000 LC pump (ThermoFisher Scientific). Fractionated peptides were separated using a 180 min gradient at 500 nL/min on a 35 cm column (i.d. 100 μm, Accucore, 2.6 μm, 150 Å) packed in-house. MS1 data were collected in the Orbitrap (120,000 resolution; maximum injection time 50 ms; AGC $10 \times 10^5$). Charge states between 2 and 5 were required for MS2 analysis, and a 180 s dynamic exclusion window was used. Top 10 MS2 scans were performed in the ion trap with CID fragmentation (isolation window 0.5 Da; Rapid;

NCE 35%; maximum injection time 35 ms; AGC $1.2 \times 10^4$). An on-line real-time search algorithm (Orbiter) was used to trigger MS3 scans for quantification.[74] MS3 scans were collected in the Orbitrap using a resolution of 50,000, NCE of 55%, maximum injection time of 200 ms, and AGC of $3.0 \times 10^5$. The close out was set at two peptides per protein per fraction.[74]

*Proteomic Data analysis*

Raw files were converted to mzXML, and monoisotopic peaks were re-assigned using Monocle.[75] Searches were performed using the Comet search algorithm against a *Rattus norvegicus* database downloaded from Uniprot in 2021. We used a 50 ppm precursor ion tolerance, 1.0005 fragment ion tolerance, and 0.4 fragment bin offset for MS2 scans collected in the ion trap, and 0.02 fragment ion tolerance; 0.00 fragment bin offset for MS2 scans collected in the Orbitrap. TMTpro on lysine residues and peptide N-termini (+304.2071 Da) and carbamidomethylation of cysteine residues (+57.0215 Da) were set as static modifications, while oxidation of methionine residues (+15.9949 Da) was set as a variable modification.

Each run was filtered separately to 1% False Discovery Rate (FDR) on the peptide-spectrum match (PSM) level. Then proteins were filtered to the target 1% FDR level across the entire combined data set. For reporter ion quantification, a 0.003 Da window around the theoretical m/z of each reporter ion was scanned, and the most intense m/z was used. Reporter ion intensities were adjusted to correct for isotopic impurities of the different TMTpro reagents according to manufacturer specifications. Peptides were filtered to include only those with a summed signal-to-noise (SN) ≥ 160 across all TMT channels. For each protein site, the filtered peptide TMTpro SN values were summed to generate protein quantification values. The signal-to-noise (S/N) measurements of peptides assigned to each protein were summed (for a given protein). These values were normalized so that the sum of the signal for all proteins in each channel was equivalent thereby accounting for equal protein loading. For the final analysis TMT channels 126, 127N, 127C, 128N, 128C, 129N, 132C, 133N, 133C, 134N, 134C, and 135N were used. Differential protein expression was then quantified using the voom function from the Limma package (version 3.56.1) to transform and normalize the expression data. Differential abundance of E3 ubiquitin ligases was performed by comparing the differential abundance to a published comprehensive list of human E3 ligases.[66] The differences in predicted half-lives of differentially abundant up vs. downregulated proteins were compared using published comprehensive dataset from mouse 3T3 fibroblasts, matched to rat proteins based on gene name. [31]

**Mouse studies**

All rodent studies were performed in accordance with NIH Guidelines on the Use of Laboratory Animals and with approval by Institutional Animal Care and Use Committees (Protocols 805309, 805255, and 807751). Mice and rats were housed in AAALAC-accredited University Laboratory Animal Resources (ULAR) animal facilities, with alternating 12-hour light-dark cycles, free access to food and water, and frequent bedding changes for all animals.

*Postnatal mouse characterization*

E18.5 mice were obtained from a timed pregnant C57BL/6J mouse. C57BL/6J wild type littermates from separate litters were sacrificed at either P2, P7, or P21. After removing the heart, either atria were trimmed off and the ventricular mass was snap frozen in liquid nitrogen or the entire heart was fixed overnight in 4% PFA followed by serial ethanol dehydration. Thiourea buffer was used to extract total protein or protein fractionation, followed by western blotting was performed as below.

*ROR2-overexpression mice*

Studies were performed in wild type C57BL/6J mice (Jackson Laboratory, Bar Harbor, ME). To generate cardiac ROR2 overexpression, AAV9 expressing HA-tagged mouse *Ror2* or HA-tagged GFP under a CMV reporter were delivered by retroorbital injection (RO) in 4-week old mice. AAV9 dosing was determined by testing 1 to 4e10 genome copies (GC)/g of *Ror2* or GFP that maximized cardiac delivery while minimizing off target expression in quadriceps, liver, and lung by western blot at 2 to 4 weeks after injection. Minimal extracardiac expression was observed in liver or lung, and RV and LV expression was 4 to 5-fold higher than in quadriceps with the highest dose at 4 weeks (**Supplement Fig. 4a**). Thus, we selected 4e10 GC/g for further studies. Mice were randomly selected from each cage to receive either AAV9-GFP or AAV9-ROR2 (2 each per cage). Mice were phenotyped and sacrificed 8-weeks after injection (age 12 weeks).

*Mouse pulmonary artery banding*

Pulmonary artery banding was performed in 9-week-old C57BL/6J mice in accordance with NIH Guidelines on the Use of Laboratory Animals and with approval by the Administrative Panel on Laboratory Animal Care in the University of Pennsylvania Cardiovascular Institute Rodent Cardiovascular Phenotyping Core. The anesthetic procedure included induction with ketamine (100mg/kg body weight) and xylazine (10mg/kg body weight) cocktail (intraperitoneal injection), tracheal intubation with a 20-guage needle, ventilation with a Harvard rodent ventilator (Harvard Apparatus, Holliston, MA) at a rate of 120-180 breaths per minute and tidal volume of 10 uL/g body weight. The main pulmonary artery trunk was dissected from surrounding tissues and constricted with a 7-0 suture tied against a 25-guage needle to generate a severe RVF. The same procedure including dissection was performed for sham mice apart from pulmonary artery ligature. Echocardiograms were performed 14 days after PAB/sham surgery as above. ROR2 knockdown in PAB mice was generated by retroorbital injection of AAV9 to deliver *Ror2*-targeting shRNA with the above sequence used for in vitro knockdown conserved in mouse/rat or scrambled shRNA at a dose of 4e10 GC/g at age 5 weeks with PAB/sham performed at 9 weeks. Mice were randomized to receive shScr vs shROR2 using GraphPad to randomly assign mice to a group for each round of injections.

*Mouse Cardiac Phenotyping*

Mouse cardiac physiology phenotyping was performed in the University of Pennsylvania Cardiovascular Institute Rodent Cardiovascular Phenotyping Core who were blinded to groups. For echocardiography, mice were sedated using Avertin by intraperitoneal injection and their temperature was maintained on a biofeedback head-pad. Mice were imaged using a MS400 18-38 MHz probe with a Vevo2100 ultrasound. Echocardiography data was analyzed using Vevo Lab software (Visualsonics). RV end diastolic pressure (RVEDP) was measured by advancing a Millar SPR-1000 Mikro-Trip Mouse Pressure Catheter (Millar, Houston Texas) via the right jugular vein until a ventricular wave form was identified. Data was analyzed using LabChart software (AD Instruments, Colorado Springs, CO).

*In vivo puromycin labeling*

Mice were singly caged just prior to puromycin injection. Mice were injected intraperitoneal with puromycin (40 nmol/g body weight diluted in PBS) using a staggered schedule to allow sufficient time for sacrifice and heart collection to ensure equal periods of puromycin labeling.[21] After 60 ± 1 minute, mice were euthanized and the hearts were dissected as above, snap frozen, and processed for protein extraction. Puromycin labeling was normalized between $ROR2^{OE}$/GFP, PAB/Sham, and $PAB^{shRor2}$/$PAB^{shScr}$ for each staggered set of injections.

*Mouse heart collection*

After euthanasia, total heart mass was assessed after trimming great vessels, and then normalized to tibia length measured with digital calipers. RV and LV mass was assessed after trimming the atria and separating the RV free wall from the septum and LV. For histology, either a 3mm-punch biopsy was collected from the RV and LV free walls, the base of the heart was isolated, or the intact heart was collected and fixed overnight in 4% PFA followed by serial ethanol dehydration. The remaining heart was separated into RV or LV components and snap frozen in liquid nitrogen. Snap frozen tissue was powdered using a liquid nitrogen cooled cryomill (Retsch).

*Mouse protein extraction*

Protein extraction was performed from snap frozen tissue using a fractionation protocol to separate cytoplasmic from the nuclear/cytoskeleton fractions.[68] Powdered snap frozen tissue was first exposed to a gentle extraction method using 0.5% Triton-X 100 in Rigor buffer (composition: 10mM Tris, 2mM EGTA, 75mM KCl, pH to 7.1) supplemented with protease and phosphatase inhibitors.[68] The tissue was incubated on ice in 20:1 (µL:mg) triton buffer for 30 minutes with inversion and flicking every 3 minutes until all samples were pale. The samples were centrifuged at 5,000G x10 min at 4°C, the supernatant contained cytoplasmic proteins while the nuclear and cytoskeleton protein was retained in the pellet. The supernatant, hereafter referred to as the nuclear-cytoskeletal depleted (“NCD”) fraction was collected and frozen. The pellet was washed in triton buffer and re-centrifuged. This second supernatant was discarded. The remaining pellet was then exposed to a harsher extraction method using TU-buffer (30:1µL:mg of original sample mass) in a Potter-Elvehjem homogenizer to solubilize all remaining proteins. This latter fraction was enriched for nuclear and cytoskeletal proteins hereafter referred to as the nuclear-cytoskeletal enriched (“NCE”) fraction (**Supplement Fig 1e**).

*Mouse cardiac histology*

Mouse histology was performed in the Penn CVI Mouse Histology core using antigen retrieval and antibodies listed in **Supplementary Table 13**. For assessment of ubiquitin staining and peripheral beta catenin, microscopy was performed using a representative section on Keyence microscope at 60X magnification. Quantification of HA-tag positive cardiomyocytes for ubiquitin staining and β-catenin localization in longitudinal orientation was performed using FIJI as above for NRVM analyses. RV cardiomyocyte cross-sectional area was quantified using WGA and the HeartJ macro to quantify the minimum Feret Diameter as a metric that is less susceptible to variability in cardiomyocyte orientation in cross-section for an average of ~500 cardiomyocytes/mouse.[67] Total RV area was quantified by imaging the entire heart in a 4-chamber view and then analyzed in FIJI. The epicardial and endocardial borders were defined using colour deconvolution to create a binary mask of the RV and then quantifying this area. For Trichrome and Sirius red percent quantification, a similar method was used to quantify only the area staining positively for fibrosis and normalizing to the total area. RNAscope was performed according to manufacturer instructions using the RNAscope® Multiplex Fluorescent V2 kit (ACD, Newark, CA) and probes to *Ror2*, *Ryr2* to identify cardiomyocytes, and *Dcn* to identify fibroblasts. DAPI was used to identify nuclei. RNAscope images were captured using Zeiss Airyscan 880 Confocal Microscope.

*qPCR*

RNA extraction was performed using RNeasy mini kits (Qiagen Hilden, Germany) according to manufacturer instructions. cDNA was made using Applied Biosystems™ High-Capacity cDNA Reverse Transcription Kit (FisherScientific). Gene expression was quantified suing SYBR Green Master Mix RT-PCR. Primer sequences (**Supplementary Table 14**).

*Western Blot*

Equal amounts of total protein or protein fractions were separated using SDS-page in a 4-20% gradient gel and then transferred to PVDF membrane. Total protein as a loading control was measured using ponceau S stain imaged using ImageQuant™ LAS 4000. Blots were blocked in 3% BSA diluted in PBS for 1 hour at room temperature (RT), incubated overnight in primary antibody, washed in TBS-T for 10 minutes x3, incubated in secondary antibody for 1 hour at RT, and washed in TBS-T for 10 minutes x3. Densitometry was assessed using SuperSignal™ West Femto enhanced chemiluminescent substrate (ThermoFisher Scientific), imaged, and quantified as previously described.[16] Primary and secondary antibodies are listed in **Supplementary Table 13**. Quantification was performed using ImageStudio (LI-COR Biotechnology, Lincoln, Nebraska) or ImageJ.

### Human Studies

Procurement of all human RV myocardial tissue was performed in accordance with protocols approved by the University of Pennsylvania Institutional Review Board (approval 802781) and the Gift-of-Life Donor Program, Philadelphia, PA. Prospective informed consent for research use of surgically explanted tissue was obtained from the transplant recipients or from the next-of-kin in the case of deceased organ donors,  as previously described.[22] RV myocardial samples were obtained from the Penn Human Heart Tissue Library collected from 2007 to 2025, including samples previously described in (Edwards, J, *et. al. Front Cardiovasc Res*. 2020).[16] RV myocardium was analyzed from patients with DCM RVF with (n=20) stratified by degree of ROR2 expression, DCM RVs with preserved RV function (n=9), and non-failing (n=11). Patient demographics and clinical data are presented in supplementary data (**Supplementary Tables 7 – 10** and **Supplementary Data 1**). Representative human samples were selected using previously published levels of ROR2 expression (n=18) as well as new samples selected to balance demographics and increase power (n=22).[16]  Proteasome capacity was measured as described above. Protein fractionation was performed for the RV tissue using Triton/TU separation as above for subsequent western blotting. Histology for RNAscope was performed similarly as for mouse histology using human specific RNAscope probes.

### Supplementary Figures and Legends

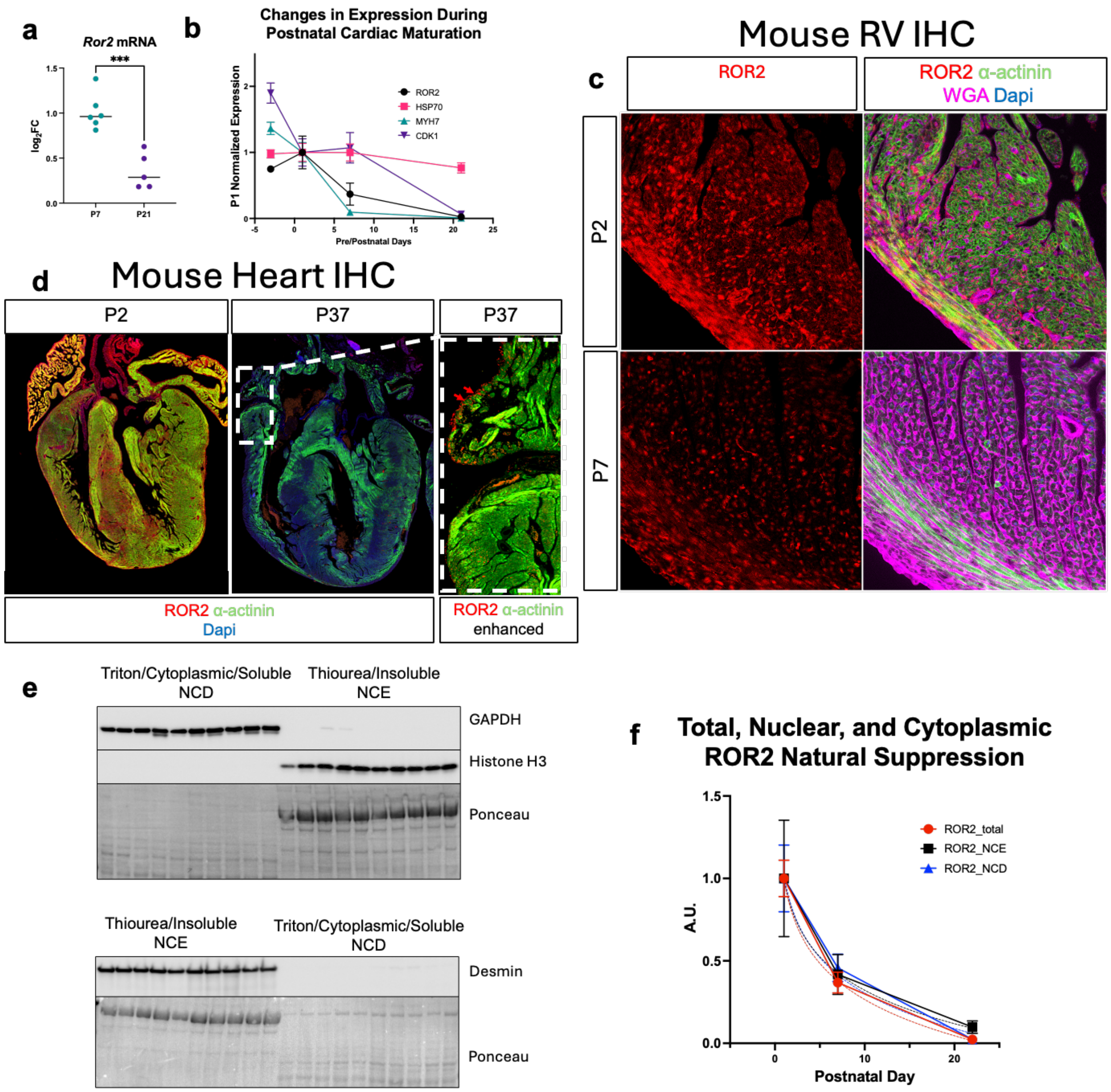

**Supplementary Figure 1 AAV9 Delivery of ROR2 to the heart and NCD vs NCE protein fractionation**

(**a**) qPCR from ventricular tissue at P7 and P21 of *Ror2* mRNA normalized to *Gapdh* and (**b**) western blot results of ROR2 protein compared to MYH7, CDK1 and HSP70 during postnatal heart maturation using thiourea extraction. (**c**) Representative immunohistochemistry of P2 or P7 RV free wall enlarged from Fig 1b and (**d**) whole heart illustrating minor amount of atrial ROR2 expression at P37 denoted by red arrows. (**e**) Protein fractionation was performed using a triton, followed by thiourea as described in the methods to separate NCE and NCD protein pools, as marked by GAPDH and Histone H3/MYH7/Desmin, respectively. (**f**) The temporal loss of total ventricular ROR2 expression was compared to NCD and NCE fractions.

Ror2 mRNA levels were measured in ventricular tissue at P7 and P21 by qPCR

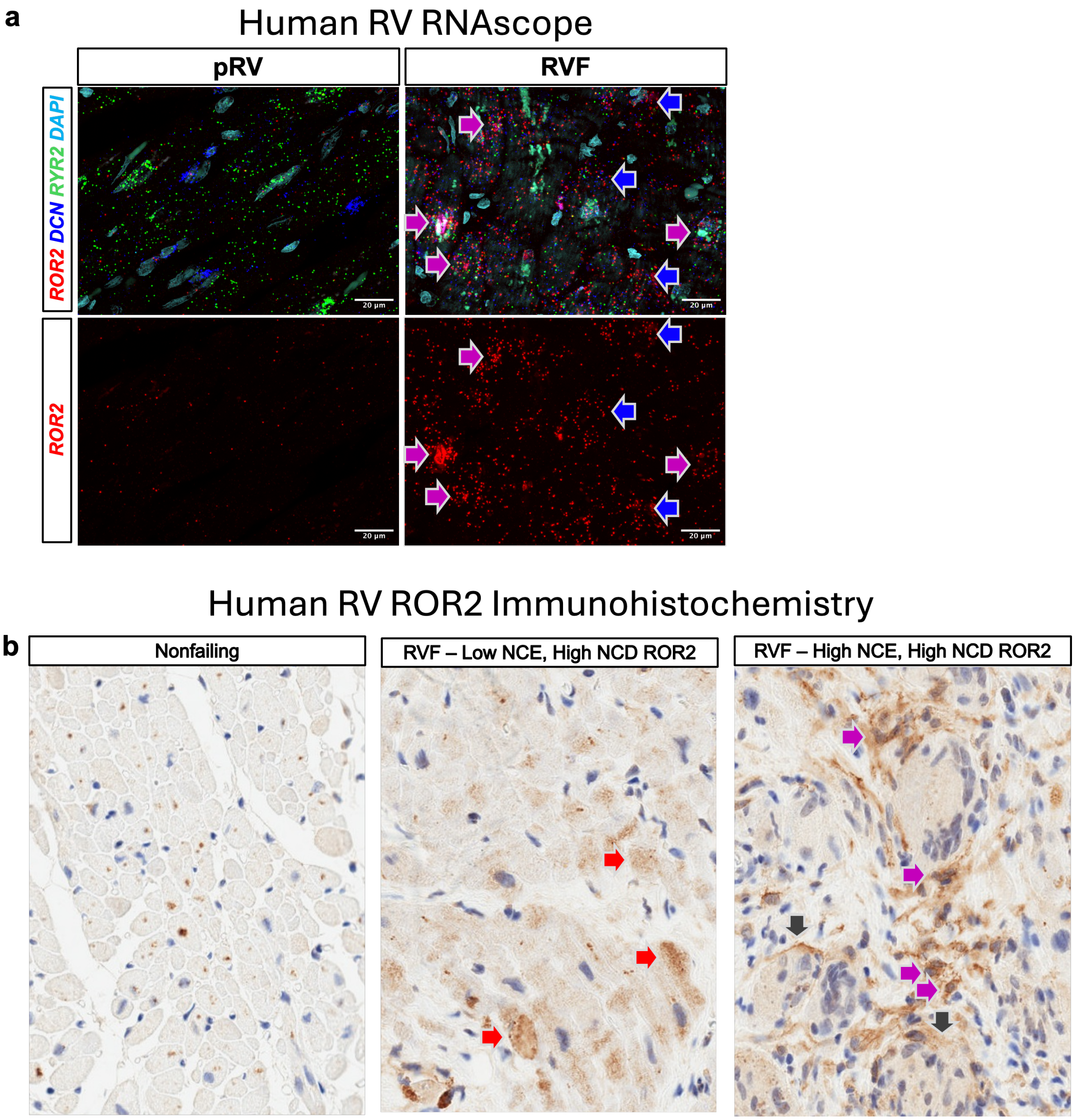

**Supplementary Figure 2 RNAscope of human RV**

**(a)** RNAscope was performed in a representative pRV (subject 1482) and RVF (subject 1656) heart tissue demonstrating robust *ROR2* expression in RVF localizing to cardiomyocytes (*RYR2,* pink arrows) and fibroblasts (*DCN,* blue arrows). (**b**) Immunohistochemistry was performed for representative nonfailing (subject 1307), RVF with low NCE/high NCD ROR2 (subject 1095), and RVF with high NCE/high NCD ROR2 (subject 1418) showing cardiomyocyte and noncardiomyocyte ROR2 localizing to cytoplasm (red arrows), cell membrane (black arrows), and nucleus ROR2 (magenta arrows).

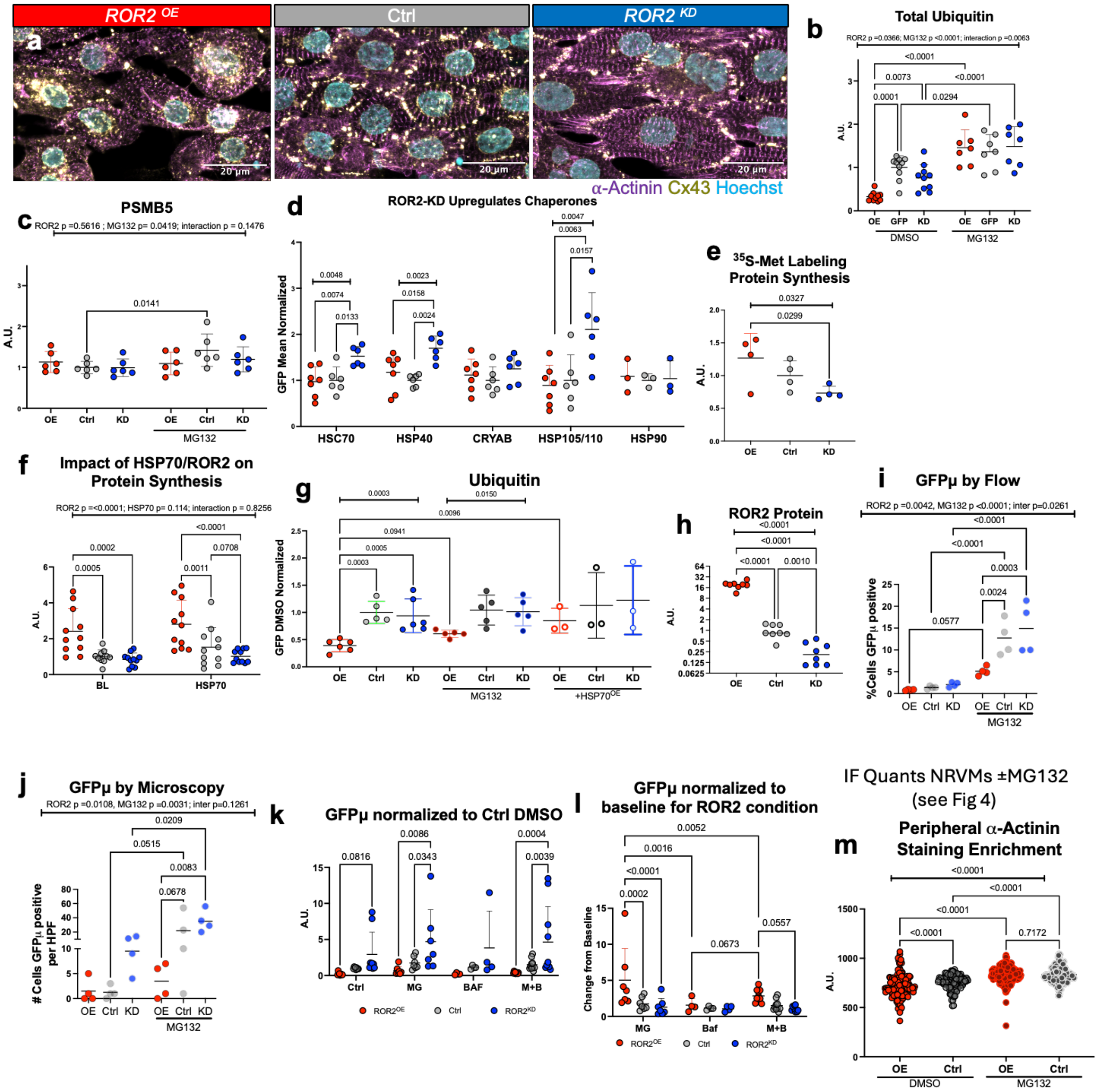

**Supplementary Figure 3 ROR2 regulates cardiomyocyte structure and proteostasis**

(**a**) Representative immunofluorescence images of ROR2$^{OE}$, Ctrl (GFP), and ROR2$^{KD}$ NRVMs stained for α-actinin and Cx43 (63X, quantifications in figure 2). (**b-d**) Western blot quantifications of total ubiquitin and PSBM5 without or with 24hours of 1µM MG132 and chaperone proteins. (**e**) Western blot quantification of $^{35}$S-Met labeled newly synthesized proteins. (**f-g**) Western blot quantification of puromycin labeled newly synthesized proteins and ubiquitin at baseline, with concomitant MG132, or with $HSP70^{OE}$ for 24 hours prior to labeling. (**h**) Quantification of ROR2 expression by western blot using non-GFP reporter adenoviruses. (**i/j**) Non-GFP reporter NRVMs were infected with adenovirus to express GFPμ for 24 hours in absence or presence of MG132 and quantified for GFPμ positivity by thresholding using microscopy and flow cytometry as described in the methods. (**k/l**) GFPμ levels were also evaluated by western blot in absence or presence of MG132, bafilomycin, or both, normalized to control+DMSO (**k**) or normalized to baseline for that specific ROR2 condition (**l**). (**m**) The impact of MG132 on $ROR2^{OE}$ induced loss of peripheral α-actinin was evaluated by microscopy as also described in figure 4.

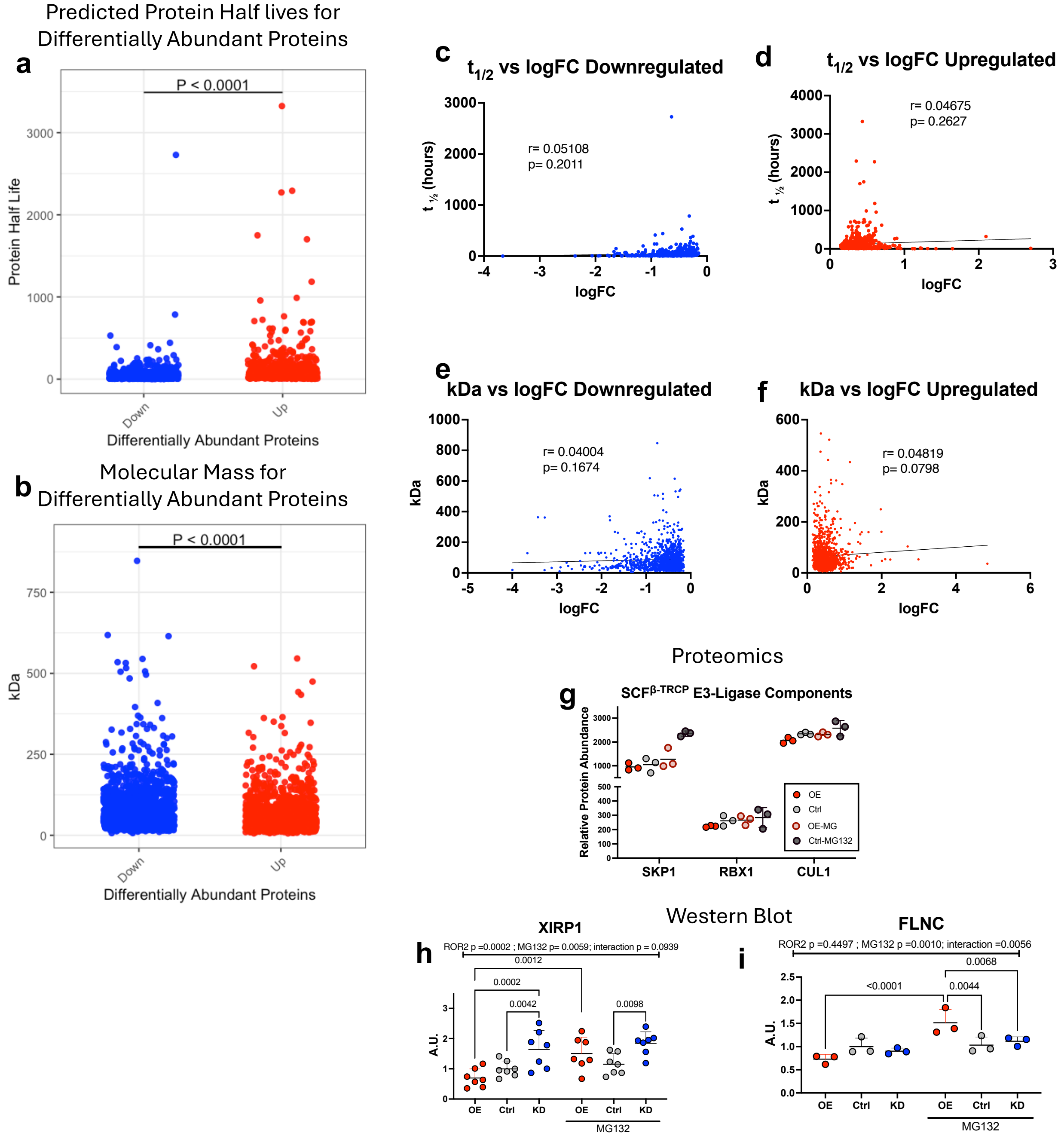

**Supplementary Figure 4. ROR2 impact on NRVM protein expression** (**a, c/d**) The predicted half-lives using published, comprehensive dataset from 3T3 mouse fibroblasts were compared for proteins that were up and down-regulated by ROR2$^{OE}$ by t-test and Pearson correlation.[31] (**b, e/f**) The molecular masses of proteins which were up and down-regulated by ROR2$^{OE}$ were compared by t-test and Pearson correlation. (**a**) Proteomic data of the components of the E3 ubiquitin ligase $SCF^{\beta\text{-TRCP}}$ that targets the transcriptional pool of β-catenin. (**h/i**) Western blot quantifications of FLNC and XIRP1 without or with 24 hours of 1μM MG132.

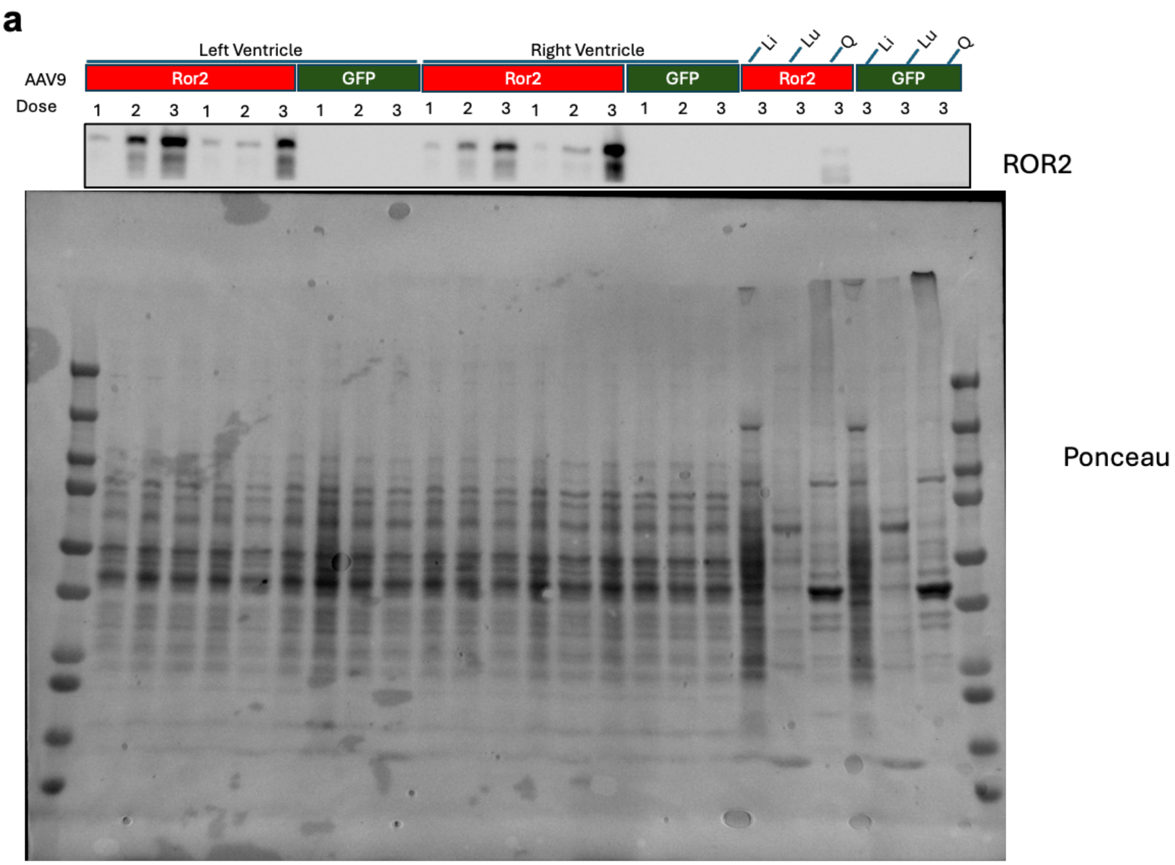

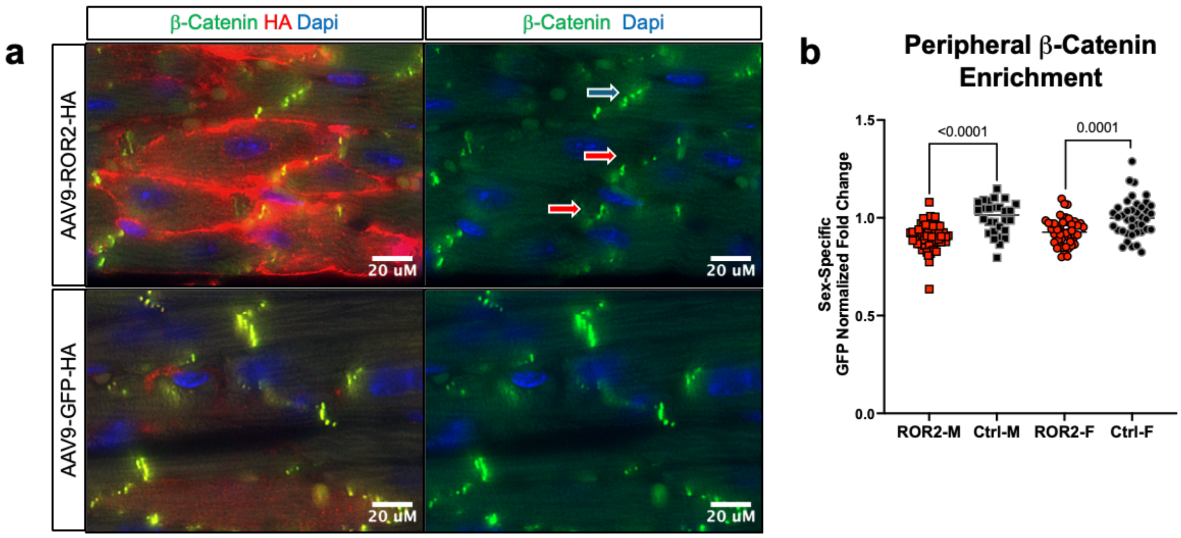

**Supplementary Figure 5 ROR2 in vivo expression by AAV9**

(**a**) Western blot for ROR2 from multiple tissues (left ventricle, right ventricle, liver (Li), lung (Lu), quadriceps (Q) four weeks after receiving low (dose 1: 1e$^{10}$ GC/g), medium (dose 2: 2e$^{10}$ GC/g) or high (dose 3: 4e$^{10}$ GC/g) AAV9 to deliver *Ror2* mRNA or *GFP*. (**b-c**) The in vivo impact of

ROR2$^{OE}$ on cell junctions was assessed by quantifying peripheral β-catenin localization in RV cardiomyocytes in vivo (60X magnification).

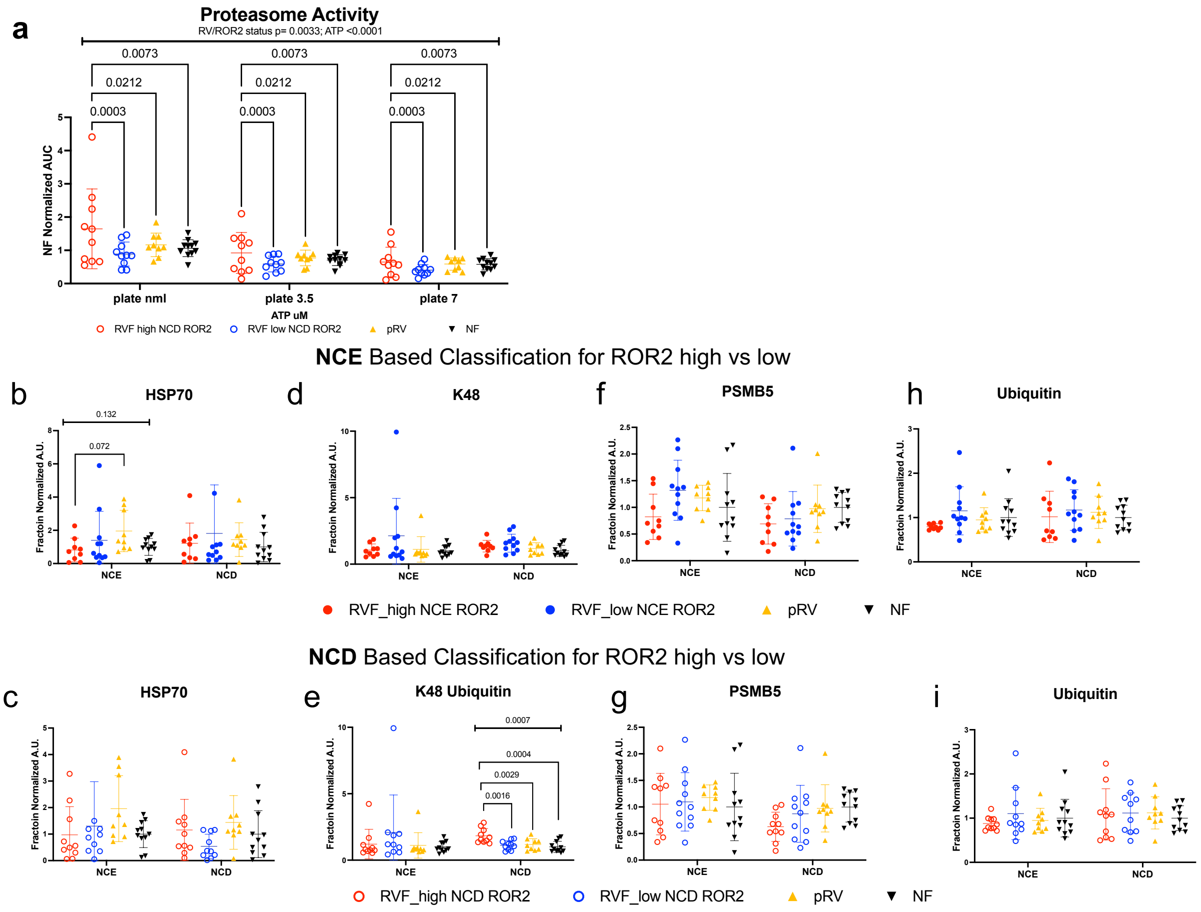

**Supplementary Figure 6 High ROR2 expression in RVF associates with markers of altered proteostasis**

Dilated cardiomyopathy human RVs were stratified by preserved RV function (pRV, orange triangles), high NCE or NCD ROR2 (red filled in or open circles) or low NCE or NCD ROR2 (blue filled in or open circles) and compared nonfailing (NF, black triangles). Proteasome capacity is increased in high NCD and NCE ROR2 (see also Fig 7). (**b-i**) pRV is associated with a trend towards higher NCE fraction HSP70 (**b**) and RVF with high NCD ROR2 demonstrates higher NCD K48 ubiquitin (**g**). No changes were observed for PSMB5 or total ubiquitin.

**Supplementary Tables**

Supplementary Table 1 – Echo parameters for PAB vs Sham mice

Supplementary Table 2 – Assessment for enrichment of Ctnnb1-responsive genes in NRVM ROR2$^{OE}$ vs Ror2$^{KD}$ RNAseq

Supplementary Table 3 – Echo and hemodynamic catheterization data analysis from AAV9 Ror2 vs GFP mice

Supplementary Table 4 – Compiled and normalized area under the curve data from AAV9 ROR2 vs GFP proteasome activity

Supplementary Table 5– Compiled and normalized area under the curve data from sham vs PAB proteasome activity

Supplementary Table 6 – Compiled echocardiographic and histologic analyses from PAB$^{shScr}$ vs PAB$^{shRor2}$

Supplementary Table 7 – Quantification of human RVF, pRV, and Nonfailing demographic and anthropometric data

Supplementary Table 8 – Quantification of human RVF and pRV demographic, anthropometric, and clinical data

Supplementary Table 9 – Quantification of human RVF with high vs low NCE ROR2 demographic, anthropometric, and clinical data

Supplementary Table 10 – Quantification of human RVF with high vs low NCD ROR2 demographic, anthropometric, and clinical data

Supplementary Table 11– Compiled and normalized area under the curve data from human RV proteasome activity

Supplementary Table 12 – ROR2 NCE and NCD expression correlation against proteasome capacity in human RVF

Supplementary Table 13 – Antibody and RNAscope probes

Supplementary Table 14 – qPCR primer sequences

1363 Supplementary Table 1

| Variable | Sham (n=6) | PAB (n=12) | P | NCE r | NCE p | NCD r | NCD p |
|---|---|---|---|---|---|---|---|
| BW | 25.62 ± 1.67 | 26.54 ± 1.72 | 0.299 | 0.005 | 0.987 | -0.168 | 0.604 |
| RV mass | 20.08 ± 5.13 | 38.53 ± 9.71 | 7.79E-05 | 0.003 | 0.992 | -0.218 | 0.494 |
| LV+S | 83.65 ± 8.64 | 68.59 ± 5.98 | 0.006 | 0.225 | 0.482 | 0.070 | 0.835 |
| TL | 16.67 ± 0.42 | 16.5 ± 0.48 | 0.465 | 0.209 | 0.514 | 0.236 | 0.457 |
| RV/LV+S | 0.24 ± 0.05 | 0.57 ± 0.18 | 3.64E-05 | -0.080 | 0.805 | -0.147 | 0.651 |
| RV/BW | 0.78 ± 0.17 | 1.45 ± 0.36 | 7.37E-05 | 0.001 | 0.998 | -0.119 | 0.716 |
| RV/TL | 1.21 ± 0.3 | 2.34 ± 0.61 | 7.58E-05 | -0.020 | 0.952 | -0.154 | 0.635 |
| BW/TL | 1.54 ± 0.1 | 1.61 ± 0.08 | 0.177 | -0.115 | 0.722 | -0.469 | 0.128 |
| LV+S/BW | 3.26 ± 0.15 | 2.59 ± 0.22 | 1.89E-06 | 0.219 | 0.494 | 0.252 | 0.430 |
| HR | 609 ± 29 | 570 ± 31 | 0.029 | -0.379 | 0.224 | -0.007 | 0.997 |
| RV Area Diastole (mm^2) | 4.28 ± 0.75 | 9.21 ± 3.42 | 3.66E-04 | 0.702 | 0.011 | -0.035 | 0.921 |
| RV Area Diastole (mm^2) /BW(g) | 0.26 ± 0.04 | 0.53 ± 0.2 | 4.30E-04 | 0.765 | 0.004 | -0.168 | 0.604 |
| RV Area Systole | 1.55 ± 0.53 | 5.21 ± 3.13 | 0.002 | 0.700 | 0.012 | 0.056 | 0.869 |
| FAC | 61.35 ± 18.26 | 47.3 ± 14 | 0.087 | -0.620 | 0.032 | -0.343 | 0.276 |
| RV Free Wall Thickness | 0.51 ± 0.16 | 0.68 ± 0.15 | 0.043 | -0.141 | 0.663 | 0.074 | 0.821 |
| TAPSE | 1.02 ± 0.15 | 0.82 ± 0.23 | 0.049 | -0.559 | 0.059 | -0.301 | 0.342 |
| TDI S Wave | 35.77 ± 8.07 | 27.15 ± 3.86 | 0.007 | -0.376 | 0.223 | -0.321 | 0.307 |
| Pk Gradient mmHg | 3.34 ± 1.27 | 66.09 ± 13.75 | 4.72E-09 | -0.111 | 0.732 | -0.028 | 0.939 |

| | | | |
|---|---|---|---|
| TR Observed ≥Mod | 0% | 42% | 0.11 |

Echo parameters for PAB vs Sham mice presented as mean ± standard deviation or percents with statistical analysis by t-test or Fisher exact. Parameters were also assessed for correlation to level of sham-normalized ROR2 expression in the NCE and NCD fractions using Pearson correlation.

Supplementary Table 2

| | Sig ROR2 | Non-sig Ror2 |
|---|---|---|
| Sig *Ctnnb1* | 43 | 212 |
| Non-sig *Ctnnb1* | 1436 | 8309 |

OR 1.174

Chi-Square P 0.3449

Enrichment of CTNNB1-responsive genes were assessed by Chi-Square using data from GSE121234.

Supplementary Table 3

| Metric | Male GFP | Male Ror2 | *P* | Female GFP | Female Ror2 | *P* |
|---|---|---|---|---|---|---|
| Echo Body wgt (g) | 27.2 ± 0.8 | 27.0 ± 1.5 | 0.794 | 22.6 ± 1.1 | 22.3 ± 2.1 | 0.82 |
| HR | 607 ± 39 | 544 ± 47 | 0.084 | 540 ± 77 | 522 ± 43 | 0.694 |
| LVPWd | 1.07 ± 0.1 | 0.93 ± 0.1 | 0.022 | 1.02 ± 0.1 | 1.00 ± 0.1 | 0.791 |
| IVSd | 1.07 ± 0.1 | 0.94 ± 0.1 | 0.08 | 1.02 ± 0.1 | 1.02 ± 0 | 0.924 |
| LVIDd | 3.18 ± 0.2 | 3.56 ± 0.1 | 0.011 | 3.28 ± 0.1 | 3.04 ± 0.1 | 0.025 |
| LVPWs | 1.39 ± 0.1 | 1.28 ± 0.1 | 0.099 | 1.28 ± 0.1 | 1.31 ± 0.1 | 0.612 |
| IVSs | 1.43 ± 0 | 1.26 ± 0.1 | 0.007 | 1.34 ± 0.1 | 1.41 ± 0 | 0.301 |
| LVIDs | 1.81 ± 0.1 | 2.39 ± 0.1 | 2.41E-04 | 2.08 ± 0.3 | 1.84 ± 0.1 | 0.196 |
| FS | 43 ± 2.4 | 33 ± 2.4 | 0.001 | 36.8 ± 7.5 | 39.4 ± 2.5 | 0.525 |
| EDV | 31.6 ± 4.6 | 43.4 ± 3.5 | 0.006 | 35.9 ± 5.6 | 32.9 ± 5 | 0.454 |
| ESV | 9.5 ± 1.4 | 17.2 ± 1.8 | 5.05E-04 | 11.6 ± 3.1 | 10.8 ± 2.5 | 0.707 |
| EF% | 69.9 ± 5.1 | 60.0 ± 3.9 | 0.022 | 67.4 ± 5.2 | 66.9 ± 4.2 | 0.885 |
| RVOT mm sys | 1.5 ± 0.2 | 1.8 ± 0.1 | 0.035 | 1.7 ± 0.1 | 1.8 ± 0.1 | 0.098 |
| RVOT mm dia | 1.7 ± 0.1 | 2.0 ± 0.1 | 0.008 | 1.9 ± 0.2 | 2.0 ± 0.2 | 0.233 |
| TAPSE | 1.1 ± 0.1 | 0.8 ± 0.1 | 0.008 | 1.1 ± 0.2 | 1.0 ± 0.1 | 0.129 |
| E | 603 ± 37 | 588 ± 70 | 0.713 | 569 ± 113 | 593 ± 63 | 0.726 |
| S' | 31.2 ± 2.2 | 26.3 ± 4.7 | 0.111 | 29.9 ± 4.4 | 30.5 ± 4.2 | 0.837 |
| E' | 30.3 ± 2.1 | 24.0 ± 2.1 | 0.005 | 30.5 ± 8 | 29.1 ± 4 | 0.761 |
| E/E' | 20 ± 2.1 | 24.5 ± 0.9 | 0.007 | 19.1 ± 2.6 | 20.8 ± 4.1 | 0.505 |
| LVEDV/BW | 1.2 ± 0.2 | 1.6 ± 0.1 | 0.008 | 1.6 ± 0.3 | 1.5 ± 0.1 | 0.426 |
| TL | 17.5 ± 0.4 | 17.2 ± 0.3 | 0.13 | 17.4 ± 0.2 | 17.4 ± 0.6 | 0.924 |
| LVEDV/TL | 1.8 ± 0.3 | 2.5 ± 0.2 | 0.005 | 2.1 ± 0.3 | 1.9 ± 0.3 | 0.454 |
| RVOT/TL | 0.095 ± 0.01 | 0.117 ± 0.01 | 0.002 | 0.108 ± 0.01 | 0.116 ± 0.01 | 0.304 |
| RVEDP (mmHg) | -0.3 ± 0.53 | 2.5 ± 1.8 | 0.024 | 2.3 ± 0.9 | 3.3 ± 2.6 | 0.513 |

Echo and hemodynamic catheterization data analysis from AAV9 Ror2 vs GFP mice. Data presented at mean ± standard deviation with statistics by t-test.

Supplementary Table 4

| AUC | AAV | Sex | ATP | Plate Normalized GFP | Sample | Plate | Overall_# for_ analysis group |
|---|---|---|---|---|---|---|---|
| 57910 | G | M | 0 | 1.783 | M_Plate1_G1_0_ATP | 1 | 1 |
| 12422 | G | M | 0 | 0.382 | M_Plate1_G2_0_ATP | 1 | 2 |
| 12148 | G | M | 0 | 0.374 | M_Plate1_G3_0_ATP | 1 | 3 |
| 47432 | G | M | 0 | 1.460 | M_Plate1_G4_0_ATP | 1 | 4 |
| 142344 | G | M | 0 | 0.797 | M_Plate2_G5_0_ATP | 2 | 5 |
| 214713 | G | M | 0 | 1.203 | M_Plate2_G6_0_ATP | 2 | 6 |
| 68524 | R | M | 0 | 2.110 | M_Plate1_R1_0_ATP | 1 | 1 |
| 56414 | R | M | 0 | 1.737 | M_Plate1_R2_0_ATP | 1 | 2 |
| 56641 | R | M | 0 | 1.744 | M_Plate1_R3_0_ATP | 1 | 3 |
| 58739 | R | M | 0 | 1.809 | M_Plate1_R4_0_ATP | 1 | 4 |
| 37748 | R | M | 0 | 1.162 | M_Plate1_R5_0_ATP | 1 | 5 |
| 21820 | R | M | 0 | 0.672 | M_Plate1_R6_0_ATP | 1 | 6 |
| 193500 | R | M | 0 | 1.084 | M_Plate2_R7_0_ATP | 2 | 7 |
| 228652 | R | M | 0 | 1.281 | M_Plate2_R8_0_ATP | 2 | 8 |
| 51744 | G | M | 3.5 | 1.593 | M_Plate1_G1_3.5_ATP | 1 | 1 |
| 19529 | G | M | 3.5 | 0.601 | M_Plate1_G2_3.5_ATP | 1 | 2 |
| 8707 | G | M | 3.5 | 0.268 | M_Plate1_G3_3.5_ATP | 1 | 3 |
| 56664 | G | M | 3.5 | 1.745 | M_Plate1_G4_3.5_ATP | 1 | 4 |
| 254164 | G | M | 3.5 | 1.424 | M_Plate2_G5_3.5_ATP | 2 | 5 |
| 375192 | G | M | 3.5 | 2.102 | M_Plate2_G6_3.5_ATP | 2 | 6 |
| 52306 | R | M | 3.5 | 1.611 | M_Plate1_R1_3.5_ATP | 1 | 1 |
| 69994 | R | M | 3.5 | 2.155 | M_Plate1_R2_3.5_ATP | 1 | 2 |
| 69668 | R | M | 3.5 | 2.145 | M_Plate1_R3_3.5_ATP | 1 | 3 |
| 77260 | R | M | 3.5 | 2.379 | M_Plate1_R4_3.5_ATP | 1 | 4 |
| 48675 | R | M | 3.5 | 1.499 | M_Plate1_R5_3.5_ATP | 1 | 5 |
| 22645 | R | M | 3.5 | 0.697 | M_Plate1_R6_3.5_ATP | 1 | 6 |
| 364333 | R | M | 3.5 | 2.041 | M_Plate2_R7_3.5_ATP | 2 | 7 |
| 550076 | R | M | 3.5 | 3.081 | M_Plate2_R8_3.5_ATP | 2 | 8 |

Compiled and normalized area under the curve data from AAV9 Ror2 vs GFP proteasome activity, data also presented in figure 6U.

1386

Supplementary Table 5

| | Set | Surg | ATP 0 | ATP 3.5 | Sham nmlized 0ATP | Sham nmlized 3.5 ATP |
|---|---|---|---|---|---|---|
| 256_4 | 1 | PAB | 172470 | 336969 | 1.398 | 2.730 |
| 262_16 | 1 | PAB | 173606 | 326566 | 1.407 | 2.646 |
| 343_3 | 1 | PAB | 123949 | 267936 | 1.004 | 2.171 |
| 256_3 | 1 | PAB | 169792 | 314556 | 1.376 | 2.549 |
| 343_20 | 1 | PAB | 178127 | 328130 | 1.443 | 2.659 |
| 262_4 | 1 | PAB | 271282 | 443361 | 2.198 | 3.593 |
| 262_8 | 1 | PAB | 164703 | 384208 | 1.150 | 2.683 |
| 256_10 | 1 | Sham | 123270 | 227827 | 0.999 | 1.846 |
| 262_5 | 1 | Sham | 123554 | 225791 | 1.001 | 1.830 |
| 262_19 | 1 | Sham | 143221 | 343074 | 1.000 | 2.395 |
| 519_7 | 2 | Sham | 278280 | 351535 | 1.450 | 1.831 |
| 519_9 | 2 | Sham | 252049 | 279839 | 1.313 | 1.458 |
| 519_12 | 2 | Sham | 101130 | 399173 | 0.527 | 2.080 |
| 519_14 | 2 | Sham | 136340 | 333454 | 0.710 | 1.737 |
| 519_6 | 2 | PAB | 302194 | 401072 | 1.574 | 2.089 |
| 519_8 | 2 | PAB | 452484 | 567188 | 2.357 | 2.955 |
| 519_11 | 2 | PAB | 235698 | 401174 | 1.228 | 2.090 |
| 519_13 | 2 | PAB | 319524 | 430338 | 1.665 | 2.242 |
| 519_15 | 2 | PAB | 113560 | 246119 | 0.592 | 1.282 |

Compiled and normalized area under the curve data from sham vs PAB proteasome activity.

Data also presented in Fig 7g.

Supplementary Table 6

| Metric | Scram | KD | |
|---|---|---|---|
| HR | 567 ± 39 | 575 ± 34 | 0.560 |
| BW | 24.8 ± 1.9 | 25.9 ± 1.8 | 0.090 |
| TL | 16.5 ± 0.3 | 16.6 ± 0.3 | 0.250 |
| Pk Gradient mmHg | 71.5 ± 15.5 | 66 ± 13 | 0.300 |
| Pk Velocity mm/sec | 4201.5 ± 489.1 | 4043.1 ± 408.5 | 0.340 |
| RV Area Diastole | 12.29 ± 4.2 | 8.16 ± 1.5 | 0.001 |
| RV Area Systole | 7.14 ± 3.5 | 4.09 ± 1.1 | 0.003 |
| RVEDA/TL | 0.75 ± 0.3 | 0.48 ± 0.1 | 0.0013 |
| RVEDA/BW | 0.5 ± 0.2 | 0.32 ± 0.1 | 0.0009 |
| RVESA/BW | 0.29 ± 0.1 | 0.16 ± 0 | 0.0022 |
| RV Free Wall Thickness | 0.6 ± 0.1 | 0.57 ± 0.1 | 0.290 |
| Rvwall/RVEDA | 0.055 ± 0.023 | 0.073 ± 0.019 | 0.030 |
| Rvwall/TL | 0.036 ± 0.005 | 0.034 ± 0.004 | 0.170 |
| RV wall/BW | 0.024 ± 0.004 | 0.022 ± 0.003 | 0.110 |
| TAPSE | 0.68 ± 0.15 | 0.79 ± 0.11 | 0.030 |
| e' | 37.7 ± 5.7 | 46.6 ± 5.6 | 0.020 |
| RV Sirius% | 16.9 ± 2 | 13.2 ± 4 | 0.050 |
| RV area by histo | 116.6 ± 22.7 | 88.4 ± 14 | 0.020 |
| TR ≥ Moderate | 67% | 14% | 0.0078 |

Compiled echocardiographic and histologic analyses from PAB with Ror2 KD vs scrambled shRNA. Data presented as mean ± standard deviation with statistics by t-test or Fisher Exact test.

Supplementary Table 7

| Clinical and Demographic Characteristics – RVF, pRV, and NF | | | | |
|---|---|---|---|---|
| Variable | RVF<br>N=20 | pRV<br>n=9 | NF<br>N=11 | p |
| Age | 49.6 ± 13 | 50.0 ± 12 | 52.7 ± 10 | 0.7796 |
| Male | 55% | 78% | 45% | 0.9181 |
| Ethnicity | | | | 0.7085 |
| Caucasian | 50% | 40% | 10% | |
| Black or African American | 63% | 25% | 13% | |
| Other | 73% | 27% | 0% | |
| Weight (kg) | 79.3 ± 17 | 94.2 ± 22 | 87.2 ± 21 | 0.1511 |
| Height (m) | 170.7 ± 9 | 176.2 ± 12 | 167.5 ± 9 | 0.1679 |
| Body surface area ($m^2$) | 1.9 ± 0.2 | 2.1 ± 0.3 | 2 ± 0.2 | 0.1331 |
| GFR | 77.5 ± 39 | 80.3 ± 19 | 93.4 ± 26 | 0.426 |

Data presented as mea

Supplementary Table 8

| Clinical and Demographic Characteristics– RVF and pRV | | | |
|---|---|---|---|
| Variables | RVF<br>n = 20 | pRV<br>n = 9 | p |
| Age | 49.6 ± 13 | 50 ± 12 | 0.9318 |
| Male | 55% | 78% | 0.4118 |
| Ethnicity | | | 0.8419 |
| Caucasian | 50% | 40% | |
| Black or African American | 63% | 25% | |
| Other | 73% | 27% | |
| Weight (kg) | 79.3 ± 17 | 94.2 ± 22 | 0.0532 |
| Body surface area ($m^2$) | 1.9 ± 0.2 | 2.1 ± 0.3 | 0.0514 |
| RA | 14.9 ± 5 | 5.3 ± 2 | <0.0001 |
| RA:PCWP | 1.2 ± 1 | 0.2 ± 0.1 | 0.0004 |
| CI | 2.0 ± 1 | 2.1 ± 0 | 0.7858 |
| Tricuspid Insufficiency >Mild | 55% | 11% | 0.0531 |
| LVEF | 18.9 ± 11 | 12.5 ± 4 | 0.1049 |
| GFR | 77.5 ± 39 | 80.3 ± 19 | 0.8211 |
| Diabetes Mellitus | 11% | 33% | 0.2872 |
| Thyroid medication | 30% | 0% | 0.1375 |
| Pacer | 45% | 56% | 0.6999 |
| ACE-i | 40% | 67% | 0.2451 |
| ARB | 15% | 33% | 0.3391 |
| β-blocker | 90% | 89% | 1 |
| Any Remodeling | 100% | 100% | 1 |
| Digoxin | 20% | 11% | 1 |
| Diuretic | 95% | 100% | 1 |
| Milrinone | 60% | 67% | 1 |

Supplementary Table 9

| Clinical and Demographic Characteristics RVF- stratified by NCE ROR2 | | | |
|---|---|---|---|
| Variables | High NCE ROR2* n = 9 | Low NCE ROR2 n = 11 | p |
| Age | 49.9 ± 17 | 49.3 ± 10.6 | 0.92 |
| Male | 44% | 64% | 0.653 |
| Ethnicity | | | 0.472 |
| Caucasian | 44% | 55% | |
| Black or African American | 55% | 27% | |
| Other | 0% | 18% | |
| Weight (kg) | 75.2 ± 13 | 82.6 ± 16.7 | 0.33 |
| Body surface area ($m^2$) | 1.87 ± 0.2 | 1.98 ± 0.3 | 0.26 |
| RA | 14.2± 3 | 15.5 ± 6 | 0.587 |
| RA:PCWP | 1.25 ± 0.8 | 1.08± 0.6 | 0.598 |
| CI | 1.91 ± 0.5 | 2.1 ± 0.8 | 0.497 |
| Tricuspid Insufficiency >Mild | 67% | 45% | 0.406 |
| LVEF | 18.9 ± 14 | 18.9 ± 8 | 0.996 |
| GFR | 55.8 ± 40 | 65.7 ± 42 | 0.0318 |
| Diabetes Mellitus | 0% | 18% | 0.479 |
| Thyroid medication | 11% | 45% | 0.157 |
| Pacer | 44% | 45% | 1 |
| ACE-i | 44% | 36% | 1 |
| ARB | 11% | 18% | 1 |
| β-blocker | 78% | 100% | 0.190 |
| Any Remodeling | 100% | 100% | 1 |
| Digoxin | 33% | 9% | 0.285 |
| Diuretic | 89% | 100% | 0.450 |
| Milrinone | 67% | 55% | 0.667 |

Data presented as mean (range) or percents. P by Chi-Square test for categorical data and t-test for continuous variables. *RVF high ROR2 groups includes the patient with high ROR2 in NCD but not NCE fraction.

Supplementary Table 10

| Clinical and Demographic Characteristics RVF- stratified by NCD ROR2 | | | |
|---|---|---|---|
| Variables | High ROR2* n = 10 | Low ROR2 n = 10 | p |
| Age | 43.5 ± 15 | 55.6 ± 8 | 0.040 |
| Male | 60% | 50% | 1 |
| Ethnicity | | | 0.0971 |
| Caucasian | 40% | 60% | |
| Black or African American | 60% | 20% | |
| Other | 0% | 20% | |
| Weight (kg) | 77.5 ±16 | 81.1 ± 17 | 0.637 |
| Body surface area ($m^2$) | 1.91± 0.2 | 1.95 ± 0.2 | 0.75 |
| RA | 14.2± 3 | 15.6± 6 | 0.521 |
| RA:PCWP | 1.50 ± 0.8 | 0.822± 0.3 | 0.021 |
| CI | 1.87 ± 0.7 | 2.18 ± 0.6 | 0.291 |
| Tricuspid Insufficiency >Mild | 40% | 70% | 0.370 |
| LVEF | 17.3 ± 14 | 20.5 ± 8 | 0.525 |
| GFR | 87.1 ± 43 | 68.0 ± 34 | 0.314 |
| Diabetes Mellitus | 20% | 0% | 0.474 |
| Thyroid medication | 20% | 40% | 0.629 |
| Pacer | 40% | 50% | 1 |
| ACE-i | 60% | 20% | 0.170 |
| ARB | 0% | 30% | 0.211 |
| β-blocker | 80% | 100% | 0.474 |
| Any Remodeling | 100% | 100% | 1 |
| Digoxin | 20% | 20% | 1 |
| Diuretic | 90% | 100% | 1 |
| Milrinone | 50% | 70% | 0.650 |

Supplementary Table 11

| | | NF 0 ATP average Normalized AUC | | | ROR2 Expression | |
|---|---|---|---|---|---|---|
| | | ATP | | | | |
| ID | Group | 0 | 3.5 | 7 | ROR2 NCE | ROR2 NCD |
| Set 1/2 | | | | | | |
| P01095 | RVF | 0.672 | 1.361 | 1.188 | 0.53 | 3.32 |
| P01149 | RVF | 0.836 | 0.502 | 0.417 | 0.94 | 0.73 |
| P01163 | RVF | 1.122 | 0.635 | 0.471 | 0.76 | 1.61 |
| P01298 | RVF | 2.241 | 1.220 | 0.555 | 3.88 | 2.51 |
| P01307 | NF | 0.555 | 0.358 | 0.281 | 0.63 | 0.63 |
| P01309 | RVF | 1.458 | 0.854 | 0.729 | 1.23 | 1.09 |
| P01418 | RVF | 4.414 | 2.098 | 1.546 | 7.27 | 2.27 |
| P01466 | pRV | 1.153 | 0.768 | 0.689 | 1.36 | 1.14 |
| P01472 | pRV | 1.408 | 0.889 | 0.756 | 0.74 | 0.75 |
| P01497 | NF | 1.139 | 0.737 | 0.613 | 1.42 | 1.12 |
| P01537 | RVF | 0.945 | 0.566 | 0.423 | 1.45 | 1.19 |
| P01538 | RVF | 0.616 | 0.382 | 0.317 | 1.64 | 1.80 |
| P01539 | NF | 0.791 | 0.442 | 0.415 | 0.97 | 1.14 |
| P01561 | NF | 0.975 | 0.578 | 0.561 | 0.95 | 1.24 |
| P01563 | pRV | 1.414 | 0.874 | 0.764 | 0.59 | 2.08 |
| P01581 | NF | 1.234 | 0.651 | 0.697 | 0.84 | 0.95 |
| P01582 | NF | 1.307 | 0.843 | 0.698 | 1.19 | 0.91 |
| P01656 | RVF | 1.710 | 0.454 | 0.546 | 2.40 | 2.96 |
| Set 3/4 | | | | | | |
| P01640 | pRV | 0.662 | 0.416 | 0.340 | 1.216 | 1.392 |
| P01643 | RVF | 0.739 | 0.136 | 0.106 | 2.131 | 5.520 |
| P01694 | RVF | 0.471 | 0.422 | 0.394 | 3.176 | 2.709 |
| P01709 | RVF | 0.406 | 0.320 | 0.265 | 0.228 | 1.108 |
| P01718 | NF | 0.863 | 0.583 | 0.419 | 1.114 | 1.397 |
| P01742 | RVF | 1.395 | 0.890 | 0.604 | 1.279 | 1.816 |
| P01745 | NF | 1.137 | 0.922 | 0.743 | 1.071 | 0.971 |
| P01756 | RVF | 2.590 | 1.370 | 0.657 | 1.383 | 2.185 |
| P01776 | pRV | 1.839 | 1.201 | 0.804 | 0.819 | 2.015 |
| P01784 | RVF | 1.641 | 1.141 | 0.781 | 1.157 | 2.561 |
| P01803 | RVF | 0.550 | 0.351 | 0.192 | 0.514 | 2.883 |
| P01807 | RVF | 0.853 | 0.627 | 0.308 | 0.481 | 1.598 |
| P01821 | pRV | 1.087 | 0.730 | 0.418 | 0.579 | 1.035 |
| P01828 | NF | 0.908 | 0.686 | 0.437 | 0.426 | 0.906 |
| P01837 | pRV | 1.092 | 0.756 | 0.500 | 0.166 | 1.693 |
| P01848 | RVF | 0.665 | 0.405 | 0.266 | 0.614 | 2.214 |
| P01883 | pRV | 1.058 | 0.843 | 0.665 | 0.936 | 0.849 |
| P01898 | pRV | 0.775 | 0.455 | 0.350 | 0.826 | 1.523 |
| P01920 | RVF | 0.410 | 0.218 | 0.153 | 1.298 | 1.299 |
| P01927 | NF | 1.120 | 0.873 | 0.625 | 1.345 | 0.707 |

| | | | | | | |
|---|---|---|---|---|---|---|
| P01936 | NF | 0.972 | 0.704 | 0.500 | 1.043 | 1.018 |
| P01958 | RVF | 0.828 | 0.883 | 0.364 | 2.140 | 0.765 |

Compiled and normalized area under the curve data from human RV proteasome activity and ROR2 expression in NCE and NCD fractions normalized to 0ATP nonfailing. High NCE and NCD RVF ROR2 samples noted by yellow highlighting.

Supplementary Table 12

| | ROR2 NCE vs. ATP_0 | ROR2 NCE vs. ATP_3.5 | ROR2 NCE vs. ATP_7 |
|---|---|---|---|
| NCE ROR2 | | | |
| Pearson r | 0.8169 | 0.6849 | 0.6883 |
| P | <0.0001 | 0.0009 | 0.0008 |
| NCD ROR2 | | | |
| Pearson r | 0.05949 | -0.06127 | 0.02961 |
| P | 0.8033 | 0.7975 | 0.9014 |

Pearson correlations between NCE and NCD ROR2 expression with proteasome capacity in human DCM RV also presented in figure 8.

Supplementary Table 13

| Target | Company | Cat Number |
|---|---|---|
| | Antibodies | |
| Alpha actinin[$] | Abcam | ab9465 |
| Beta catenin[$] | CST | 8480 |
| CDK1 | Abcam | ab32094 |
| CRYAB | Abcam | 13497 |
| Cx43[$] | Abcam | ab113700 |
| FLNC | Novus | NBP1-89300 |
| GAPDH | CST | 5174 |
| GFP | CST | 2956S |
| HA Tag | CST | 3724 |
| HISTONE H3 | CST | 4499 |
| HSC70 | Novus Bio | NBP1-97868 |
| HSP105/110 | Abcam | AB109624 |
| HSP40 | CST | 4868S |
| HSP70 | CST | 4872 |
| HSP90 | CST | 4874 |
| K48 Ubiquitin | CST | 8081 |
| K-63 Ubiquitin | CST | 5621 |
| MYH7 | Sigma | m8421 |
| PSMB5 | CST | 12919s |
| Puromycin | Millipore Sigma | MABE343 |
| ROR2 | CST | 88639 |
| ROR2[*] | Abcam | ab309483 |
| Ubiquitin | CST | 43124 |
| Ubiquitin[*] | Cytoskeleton, Inc | AUB01 |
| XIRP1 | Santa Cruz | sc-166658 |
| | RNAscope Probes - Mouse | |
| *Dcn* | ACD Bio | 413281-C3 |
| *Ror2* | ACD Bio | 430041-C2 |
| *Ryr2* | ACD Bio | 479981 |
| | RNAscope Probes - Human | |
| *DCN* | ACD Bio | 589521-C4 |
| *ROR2* | ACD Bio | 408601 |
| *RYR2* | ACD Bio | 415831-C2 |

Antibodies used for western blot and immunohistochemistry and RNAscope probes. Where a target was assessed by immunohistochemistry (IHC)/Immunofluorescence (IF), [*]denotes that the antibody was used for only immunofluorescence/immunohistochemsitry and [$]denotes an antibody that was also used for western blot.

Supplementary Table 14

| Target | Species | Fwd | Reverse |
|---|---|---|---|
| *Hspa1b* | Mus musculus | GGTGAACTACAAGGGCGAGA | CTTCATCTTCGTCAGCACCA |
| *Nppb* | Mus musculus | TTTGGGCTGTAACGCACTGA | ACTTCAAAGGTGGTCCCAGAG |
| *Gapdh* | Mus musculus | AGGTCGGTGTGAACGGATTTG | TGTAGACCATGTAGTTGAGGTCA |

qPCR primer sequences

**Supplementary and Source Data**

SD1 – Compiled clinical and demographic data for human heart tissue samples

SD2 – Full echo data and tissue weights for PAB vs Sham

SD3 – Compiled data from NRVM immunofluorescent microscopy quantifications

SD4 – Full differential gene expression data for NRVMs ROR2$^{OE}$ vs ROR2$^{KD}$

SD5 – Compiled differential gene expression from GEO GSE121234 RNAseq data from Ctnnb1 gain of function vs control compared to our RNAseq differential gene expression data for NRVMs ROR2$^{OE}$ vs ROR2$^{KD}$

SD6 - Compiled data from NRVM immunofluorescent microscopy quantifications with and without MG132

SD7 - Compiled data from NRVM immunofluorescent microscopy quantifications with and without Hspa1b overexpression

SD8 - Compiled data from NRVM pacing quantifications with and without Hspa1b overexpression

SD9 – Raw tracing data for contraction graph presented in figure

SD10 – Raw fluorescent data for NRVM proteasome activity assay

SD11 – Luciferase refolding data for luciferase activity

SD12 – Pathway enrichment analysis results for NRVM RNAseq upregulated genes in $ROR2^{OE}$

SD13– Pathway enrichment analysis results for NRVM RNAseq downregulated genes in $ROR2^{OE}$

SD14 – Full differential protein abundance for $ROR2^{OE}$ vs ctrl

SD15 -- Full differential protein abundance for $ROR2^{OE}$ DMSO vs MG132

SD16 -- Full differential protein abundance for MG132 treated $ROR2^{OE}$ vs ctrl

SD17 – Pathway enrichment analysis for proteomics $ROR2^{OE}$ vs ctrl – upregulated

SD18 – Pathway enrichment analysis for proteomics $ROR2^{OE}$ vs ctrl – downregulated

SD19 -- Pathway enrichment analysis for proteomics $ROR2^{OE}$ vs ctrl downregulated, but upregulated in $ROR2^{OE}$ vs ctrl after MG132

SD20 – Proteomics arbitrary units for E3 ubiquitin ligases as depicted in figure 4E

SD21 – Full list of E3 ligases with differential abundance data for NRVM proteomics $ROR2^{OE}$ vs ctrl and $ROR2^{OE}$ DMSO vs MG132

SD22 – Comparison of molecular mass and predicted half-lives for proteins that are up- vs down-regulated in ROR2$^{OE}$ NRVMs.

SD23 – Echo and catheterization data for Ror2 vs GFP overexpression in vivo with AAV9

SD24 -- Source data for β-catenin peripheral localization after AAV9 ROR2 vs GFP in RV cardiomyocytes (Supp Fig 6)

SD25 – Source data for ubiquitin staining after AAV9 ROR2 vs GFP in RV cardiomyocytes (Fig 6S)

SD26 – Minimum Feret diameter for AAV9 ROR2 vs GFP in RV cardiomyocytes (Fig 6G)

SD27 – Proteasome activity raw data for AAV9 ROR2 vs GFP in RV male mice (Fig 6U)

SD28 -- Proteasome activity raw data for PAB vs Sham (Fig 7E)

SD29 – Compiled echo data for PAB$^{shScr}$ vs PAB$^{shRor2}$

SD30 -- Proteasome activity raw data for PAB$^{shScr}$ vs PAB$^{shRor2}$ (Fig 7S)

SD31 -- Proteasome activity raw data for human RV

**Additional source data**

Raw western blot quantifications as noted in excel tab description linking it to the figure panel.
